# Ensembles of Graph Attention Networks Supervised by Genotype-to-Phenotype Structures Improved Genomic Prediction Performance

**DOI:** 10.64898/2025.12.21.695855

**Authors:** Shunichiro Tomura, Owen Powell, Melanie J. Wilkinson, Mark Cooper

## Abstract

Accurate selection of favourable crop genotypes has motivated the exploration of diverse prediction algorithms for crop breeding applications. One genomic prediction method that has not been fully explored is graph attention networks (GAT). By directly analysing graphical data with the attention mechanism, GAT can incorporate the genotype-to-phenotype (G2P) structure to regularise predictions. As one potential G2P structure, a gene network can be inferred from interpretable machine learning models to effectively learn key features of prediction patterns, potentially improving prediction performance. Here, we investigated whether incorporating such data-driven prior knowledge into GAT improved prediction performance compared to GAT models representing a continuum of G2P structures, ranging from infinitesimal to fully connected. Applying the Diversity Prediction Theorem, we also combined these diverse G2P structures into an ensemble of GAT genomic prediction models to integrate complementary strengths of multiple models. The results for flowering time traits in two maize nested association mapping datasets showed a lack of consistent performance improvement in the data-driven prior knowledge GAT model. However, consistent outperformance was observed for the ensemble of GAT models. Improved predictions from the ensemble model may be driven by its ability to capture a more complete representation of the inferred gene network through the integration of information from diverse G2P structures. The observed results using the GAT methodology provided the foundation for potential performance improvement using GAT by integrating biological prior knowledge derived from omics data and empirically verified gene interactions in future research, thereby potentially enhancing the GAT ensemble performance.

## Introduction

Improving the accuracy of genomic prediction models (Meuwissen et al., 2001) has been a primary research target in crop breeding, motivated by the need for rapid and precise selection of heritable variation to accelerate genetic gain (Voss-Fels et al., 2019; Escamilla et al., 2025). This research target has promoted the continuous exploration of novel algorithms that can be applied to the datasets generated by crop breeding programs. From the early stages of genomic prediction research, parametric approaches, such as genomic best linear unbiased prediction (GBLUP) and Bayesian regression models, have been the primary target of investigations (Azodi et al., 2019). While the parametric models are still widely employed in crop breeding today, the emergence of non-parametric models, such as machine learning and deep learning models, has diversified genomic prediction models, aiming to capture the complex genotype-by-genotype interactions underlying genomic markers (Cooper et al., 2005, 2025; Hammer et al., 2006; Crossa et al., 2025; Messina et al., 2025).

Applications of graph neural networks (Scarselli et al., 2008) provide opportunities to emulate the networks of genes that contribute to the genetic architecture of traits (Cooper et al., 2005). The algorithmic and technical advancement in deep learning models has enabled the analysis of graphically structured datasets using graph neural networks (Bhatti et al., 2023; Vatter et al., 2023). One considerable advantage of graph neural networks is the direct incorporation of topology connecting nodes (storing a collection of feature values) grounded in underlying relationships (Scarselli et al., 2008). In particular, graph attention network (GAT; Veličković et al., 2018) models have been a widely utilised graph neural network model due to the flexibility of the attention mechanism, enhancing performance by allocating higher weights to critical predictive information (Brody et al., 2021). GAT has improved prediction performance in various tasks and hence has been utilised as a benchmark model for the evaluation of new prediction models. Graph neural networks, including GAT, have already been applied to diverse research areas in biology, including drug discovery (Abate et al., 2023; Wang et al., 2025a), gene sequence analysis (Wang and Zhang, 2021; Wang et al., 2023) and disease discovery (Hernández-Lorenzo et al., 2022; Duan et al., 2023). While graph neural networks have been employed in several cases in crop breeding, such as crop yield prediction by converting the information from farming plots into nodes (Gupta and Singh, 2023) and suitability evaluation for crops (Zhang et al., 2022), the possibility of using graph neural networks for genomic prediction remains largely unexplored.

In genomic prediction for crop breeding, graph neural networks can predict target traits while considering the possible relationship between genomic markers and target phenotypes represented as the genotype-to-phenotype (G2P) structure (Cooper et al., 2005). One possible representation of a G2P structure can be a gene network inferred from a statistical or interpretable machine learning model, considering the success of data-driven graph neural network approaches in other research areas (Li et al., 2025; Tang et al., 2025; Gouvêa et al., 2026). Data-driven prior knowledge can be utilised for genomic prediction when evidence indicates traits are regulated by complex genomic marker-by-marker interactions (Cooper et al., 2005; Powell et al., 2022). The direct relationship between genomic markers and traits can represent additive effects, whereas the interactions between the genomic markers can be used to represent non-additive effects. Such data-driven prior knowledge can provide prediction models with problem-specific information, expected to guide the prediction models to efficiently capture key features that will have the greatest influence on prediction outcomes (Von Rueden et al., 2021). The benefit of such prior knowledge inclusion is especially emphasised when the training set size is small (Von Rueden et al., 2021), which is true for many crop breeding programs that can only collect a small scale of phenotype data (Ramstein et al., 2019; Mascher et al., 2024). Consequently, the integration of data-driven gene network information as prior knowledge into genomic prediction models may help capture key additive and non-additive effects, expected to improve prediction performance (Cooper et al., 2005; Zhao et al., 2025).

While graph neural networks with data-driven prior knowledge can be leveraged to improve prediction performance, other possible G2P structures have already been proposed to represent the relationship between genomic markers and traits. For instance, infinitesimal structures assume no explicit interactions among genomic markers, sharing the conceptual similarities with the standard BLUPs and Bayesian prediction models that primarily estimate additive genomic marker effects (Meuwissen et al., 2001; Cooper et al., 2021). In contrast, fully connected graph structures may detect active non-additive interactions by allowing genomic markers to directly interact with others. This structure is inspired by the fully connected mechanism of neural networks composed of neurons and their edges connecting each other, motivated by mimicking the human brain system (Rosenblatt, 1958; Dave and Dutta, 2014). The effectiveness of the fully connected structure has been demonstrated in solving a wide range of complex prediction problems with high performance (Krogh, 2008). Prior knowledge graph structure lies between the infinitesimal and fully connected structures, expected to capture a more accurate gene network by removing non-additive interactions that hardly influence the traits. This continuum of different G2P structures (Figure 1; the infinitesimal, prior knowledge and fully connected) captures a different facet of standing genetic variation of a trait genetic architecture that is usually influenced by the mixture of additive and non-additive effects, especially for complex traits. In such cases, the prediction performance of graph neural networks based on respective G2P structures can potentially capture distinct prediction patterns. This has not explicitly been investigated through empirical genomic prediction studies for crop breeding despite its potential for influencing prediction performance.

**Figure 1:**
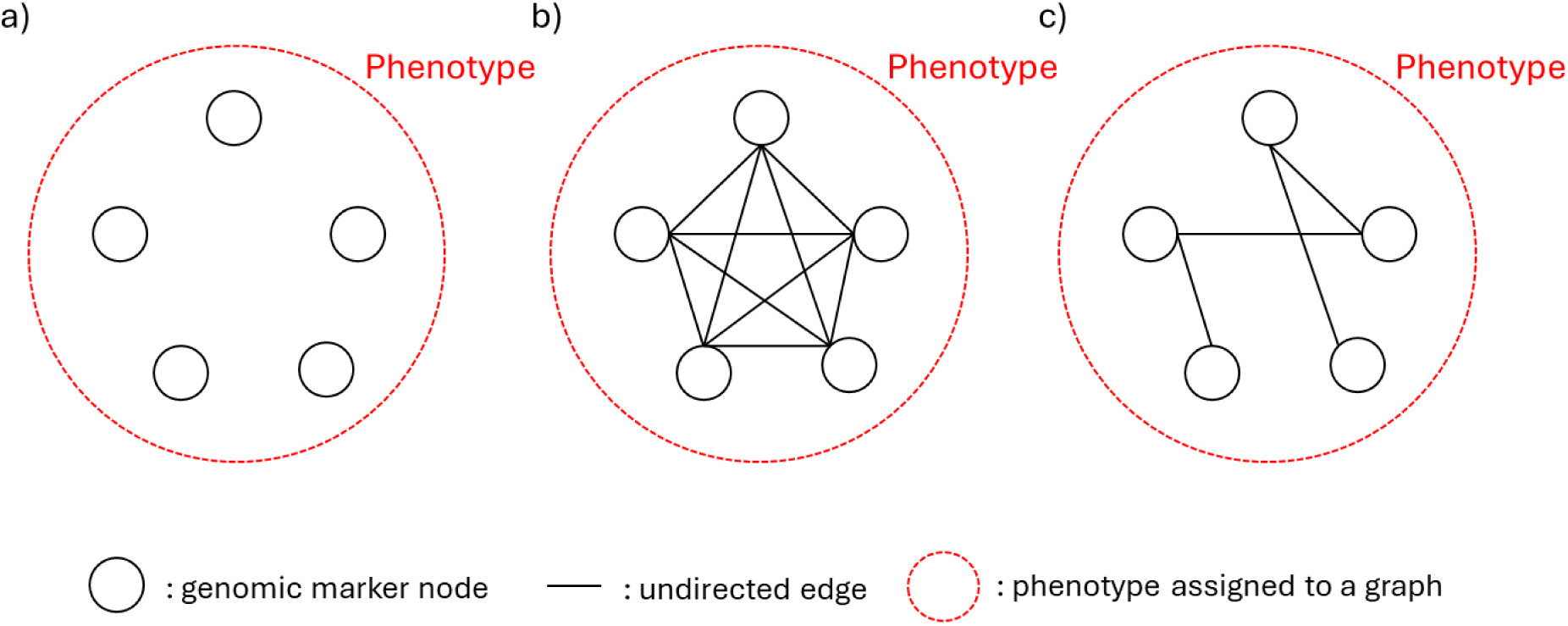
Three graph attention network (GAT) models constructed from three possible genotype-to-phenotype (G2P) relationships: a) infinitesimal model, b) fully connected model and c) data-driven prior knowledge model. The infinitesimal model assumes no explicit connections between genomic marker nodes. The fully connected model allows all genomic marker nodes to be connected to others. The data-driven prior knowledge model only allows certain connections between genomic marker nodes based on the inferred trait gene network from the provided data.

An ensemble of multiple graph neural networks grounded in different G2P structures could be another approach to improve prediction performance, as proposed by the Diversity Prediction Theorem (Page, 2018; Messina et al., 2025). The theorem suggests that an ensemble of diverse individual prediction models can be constructed to lower the prediction error and improve prediction accuracy. Experimental results have demonstrated that ensembles of multiple genomic prediction models have consistently improved prediction performance by increasing prediction accuracy and reducing prediction error in comparison to the individual genomic prediction models (Kick and Washburn, 2023; Meher et al., 2025; Tomura et al., 2025b). Hence, the ensemble of genomic prediction models, guided by diverse G2P graph structures, can also be evaluated for potential improvements beyond individual genomic prediction models.

Here, we evaluated the prediction performance of graph neural networks with three G2P graph structures (infinitesimal, fully connected and data-driven prior knowledge) and an ensemble of these graph neural networks through application to two maize experimental datasets. This study investigated the following three objectives. Firstly, the prediction performance of GAT models designed to utilise data-driven prior gene network knowledge was compared with other GAT models based on G2P graph structures that did not use the prior gene network knowledge. The prediction performance of the ensemble GAT models was also evaluated by comparing with each GAT model. Secondly, the performance decay of the GAT models with decreasing training set size was compared to analyse which GAT model is relatively less responsive to reduced training set size. Thirdly, the standing genetic variation of the trait genetic architecture captured as features by each GAT model was compared to understand the effect of diversity in genomic marker effects on the results achieved by the ensemble model.

## Materials and Methods

### 1. Datasets

We used two maize (*Zea Mays*) nested association mapping (NAM) datasets in this study, the TeoNAM (Chen et al., 2019) and MaizeNAM (Buckler et al., 2009). The TeoNAM dataset is based on samples of recombinant inbred lines (RILs) from five populations. The maize inbred line W22 was crossed with five teosinte types, four from *Z. mays ssp. parviglumis* and one from *Z. mays ssp. mexicana*. In each population, RILs were developed by controlled self-pollination following backcrossing W22 with one of the teosinte types after the initial F1 cross. All TeoNAM experiments were conducted at the University of Wisconsin West Madison Agricultural Research Station using a randomised complete block design. The TeoNAM dataset was generated by evaluating each set of RILs in two environments: the W22TIL01, W22TIL03 and W22TIL11 populations were evaluated in the summer of 2015 and 2016. The W22TIL14 population was evaluated in the summer of 2016 and 2017. The W22TIL25 population was evaluated in two environments during the 2017 summer.

The MaizeNAM dataset has 25 populations of RILs from crosses between the maize inbred B73 and 25 inbreds sampled from temperate and tropical regions. The 25 RIL populations were developed by advancing the F1 crosses to the F5 generation by controlled self-pollination. Each RIL population was evaluated in eight environments with a randomised design in the United States. Two experiments were conducted at each location in Aurora (New York), Clayton (North Carolina), Columbia (Missouri) and Urbana (Illinois). The total number of genomic markers and RILs from each dataset is summarised in Table S1.

The primary reason for selecting these two datasets is their contrast in genetic diversity. The TeoNAM dataset crossed a domesticated maize line with teosinte, an ancestral species of maize, leading to variation in allelic frequency for many genomic regions among the crosses. In contrast, the MaizeNAM contains crosses among post-domestication inbred lines developed by breeding programs, thereby driving higher levels of fixation of alleles through selective breeding in the MaizeNAM populations relative to the TeoNAM populations. Using these two NAM datasets with different levels of genomic marker diversity, we can investigate the prediction performance of the proposed approaches for a diverse range of prediction scenarios.

The two flowering time-related traits, days to anthesis (DTA) and anthesis to silking interval (ASI), were chosen for investigation in this study. For DTA, the genetic control of flowering time in maize has previously been investigated with the identification of gene pathways (Dong et al., 2012), extended by further investigation of the genetic basis for response to selection (Wisser et al., 2019). Key gene regions and regulatory pathways identified from the previous studies can be compared with the extracted trait genetic architecture from GAT models in this study. ASI is a secondary trait representing the difference between DTA and days to silking (DTS) (Leng et al., 2022), and hence the gene network might be regulated by both DTA and DTS and other mechanisms involved in the synchrony of development of the male and female flowers. This potential greater complexity in the ASI network relative to the days to flowering traits might affect the prediction performance. Using these two distinctive traits, we can investigate the predictive behaviour of the GAT and ensemble models in detail for a range of trait scenarios. For the TeoNAM dataset, the reported (Chen et al., 2019) phenotype values from two environments were used as input for the genomic prediction models. For the MaizeNAM dataset, the reported best linear unbiased predictions (BLUPs) (Buckler et al., 2009) of each RIL were used as phenotype values.

### 2. Data preprocessing

Prior to implementing the genomic prediction models, three data preprocessing methods were applied to the datasets: (1) imputing missing genomic marker alleles, (2) removing missing phenotypes and (3) reducing the total genomic markers. For imputing missing genomic marker alleles, the flanking marker imputation approach was utilised for both datasets. This approach imputes the missing allele information with the allele of the closest flanking marker. For the TeoNAM dataset, the flanking marker imputation approach is detailed in Tomura et al. (2025b) and for the MaizeNAM dataset, the approach is detailed in Buckler et al. (2009). For missing phenotypes, data points with no recorded phenotypes were removed. For the genomic marker reduction, linkage disequilibrium (LD) was employed to filter genomic markers. If LD was greater than 0.8, then one of the genomic markers was removed. The LD-filtering was implemented by PLINK (v1.9) (Chang et al., 2015) with a 30kb window and step size of 5, for both the TeoNAM and MaizeNAM datasets.

To allow for genotype-by-environment effects in the TeoNAM dataset, we combined the data from the two environments. We concatenated the two environments with a factor variable explicitly representing each environment. This enabled us to evaluate whether the genomic prediction models can predict phenotypes with higher performance using available environmental information.

### 3. Genomic prediction models

#### 3.1 Prediction mechanisms

In this study, all GAT (Veličković et al., 2018) models and interpretative analyses were implemented using the computational tool, Ensemble AnalySis with Interpretable Genomic Prediction (EasiGP; Tomura et al., 2025a), developed in Python (v3.11.10). GAT is a graph neural network that can directly analyse graphically structured data to predict target values. Graph neural networks extract key information for predictions from nodes and edges in graphs as embeddings, concentrating the information into vectors of numerical format. Hence, the extraction of embeddings containing key predictive information is a critical aspect of graph neural networks.

One significant characteristic of the GAT model is the application of a self-attention mechanism (Brody et al., 2021), assigning a weight to each piece of information from graph edges based on the level of their relevance towards prediction targets. Assigning a larger attention value to an edge passing relevant information during the embedding process helps the GAT model precisely capture key information from graphs. The attention mechanism can be written as below (Brody et al., 2021):

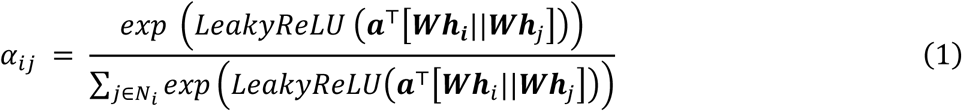

where *α*_*ij*_ is the attention value assigned to the edge between the target node *i* and a neighbouring node *j*, ***α***^⊤^ ∈ *R*^2*d*′^ is the transformed weight vector, *d*′ is the total number of attributes in each node, ***W*** is the weight matrix, *i* is the target node, *j* ∈ *N*_*i*_ is a neighbouring node of the node *i*, ***h***_*j*_ = {ℎ_1_, ℎ_2_, …, ℎ_*N*_} is the attribute set of the neighbouring nodes and || concatenates vectors. The Leaky Rectified Linear Unit (LeakyReLU; Maas et al., 2013) activation function nonlinearly converts attention values. The attention value is normalised at the end. Using attention values, predictive information is effectively extracted in each node with consideration of the information from its connected neighbouring nodes. The extracted predictive information is nonlinearly aggregated using an activation function and represented as node embeddings of each node. This calculation process can be iterated *K* times (attention layers) to stabilise the normalised attention values (multi-head attention mechanism) (Veličković et al., 2018). Node embeddings from each attention layer are concatenated to represent the final node embeddings. Summarising the predictive

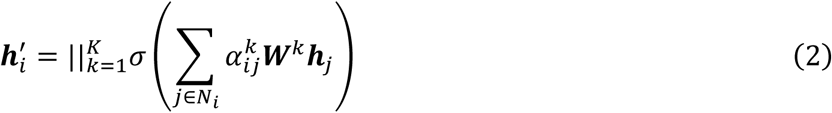

where *σ* is the nonlinear activation function and ***h***_t_′ is the updated attributes (node embeddings) of the target node *i*. These node embeddings were concatenated across nodes to calculate graph-level embeddings, which were passed to a fully connected layer to predict target values as implemented in a multi-layer perceptron.

Three distinctive GAT models (infinitesimal model, fully connected model and prior knowledge model) were developed based on three forms of possible G2P structures (Figure 1). The infinitesimal model (Figure 1a) assigned no edges between the genomic marker nodes to explicitly represent the independent contributions of the additive genomic marker effects. Thus, for the infinitesimal GAT model, no non-additive marker effects were allowed. The fully connected model (Figure 1b) predicted phenotypes using a graph with genomic marker nodes connected to all the other nodes with undirected edges. In this case, all genomic markers are assumed to interact explicitly with other genomic markers to capture the potential for non-additive marker effects between all combinations of markers. The data-driven prior knowledge model (Figure 1c) is intermediate between the infinitesimal and fully connected extremes. The implementation of the prior knowledge model considered here was supervised by a gene network inferred from a random forest (RF; Breiman, 2001) equipped with Shapley scores (Shapley, 1953) using SHapley Additive exPlanations (SHAP; Lundberg and Lee, 2017) as the data-driven prior knowledge (the calculation method is discussed in the next subsection). Shapley scores were returned for every pair of genomic markers, and the top 20% of the genomic marker interactions with the highest Shapley scores were selected to draw undirected edges as the prior knowledge graph. The extracted gene network only leverages edges from the key genomic marker-by-marker interactions identified by the Shapley scores, thereby expected to remove components of complexity and noise generated from connecting all genomic marker nodes with edges as observed in the fully connected model. The prediction performance of RF was used as a reference for the data-driven prior knowledge GAT model since both predicted phenotypes based on the same inferred marker interaction effects on the target traits. The prediction performance of the prior knowledge GAT model was expected to be higher than that of RF due to the incorporation of the attention mechanism.

Following the framework of the Diversity Prediction Theorem (Page, 2018), discussed further below, the naïve ensemble approach of Tomura et al. (2025b) was applied to the GAT models, which calculated the arithmetic mean of the predicted phenotypes from each GAT model with equivalent weight. Four distinctive naïve ensemble-average models were constructed based on all possible combinations of GAT models to compare prediction performance.

#### 3.2 GAT hyperparameter selection

Hyperparameters of the GAT models were tuned to improve the prediction performance (Table S2). The application of the same hyperparameter sets from the TeoNAM dataset did not reach high prediction performance for the MaizeNAM dataset, and hence the hyperparameter sets were tuned at the dataset level for each GAT model. For the TeoNAM dataset, one hidden layer with 50 neurons was added with a dropout rate of 0. The Exponential Linear Unit (ELU; Clevert et al., 2015) function was leveraged as an activation function for each neuron to nonlinearly aggregate extracted information in each node as node embeddings. The Adaptive Moment Estimation (Adam; Kingma, 2014) with a learning rate of 0.005 and a weight decay rate of 0.0004 was used for the optimisation. The number of heads was 1. The number of epochs was set as 200 for the infinitesimal model and 50 for the remaining models. For the MaizeNAM dataset, the same hyperparameter settings in the TeoNAM dataset were used for the number of hidden layers, neurons, activation function, optimisation function and head number. The dropout rate and epoch numbers were set as 0.5 and 200, respectively, across the GAT models. To extract key gene interactions for the prior knowledge model, RF was implemented with a total tree number of 1,000, while the default settings were used for the other hyperparameters. These selected hyperparameters were heuristically evaluated, and hence, through further investigation, other hyperparameter combinations may achieve higher prediction performance. Methods of the mechanistic hyperparameter tuning with low computational cost for graph neural networks can be investigated as future studies.

### 4. Genomic marker and marker-by-marker interaction effect estimation

From the GAT models, the genomic marker effects for target trait prediction were estimated using Integrated Gradients (Sundararajan et al., 2017) from the genomic markers as a part of the pipeline in EasiGP:

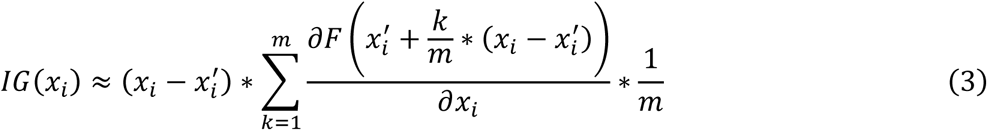

where *x* is the attribute (SNP) value, *x*^′^ is the baseline value of the attribute *x* and *m* is the total interpolation steps of the integral. Integrated Gradients measure the influence of each attribute using the cumulative values of integrals between the predicted values for individual RILs including and excluding the target attribute. Integrated Gradients calculates genomic marker effects element-wise, and thus the final genomic marker effects were calculated by summing the genomic marker effects of 50 randomly chosen individuals (RILs) from the test set. The mean values of genomic marker effects from each prediction scenario were calculated as the final marker effects in the end.

For RF, the impurity-based importance of each genomic marker was used as the genomic marker effects. The pairwise Shapley score (Shapley, 1953; Lundberg and Lee, 2017) was calculated to estimate the genomic marker-by-marker interaction effects:

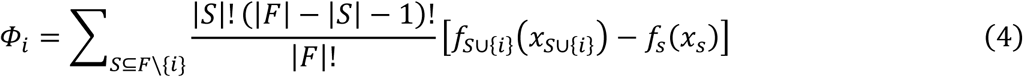

where 𝛷_*i*_ is the Shapley score of the target attribute (SNP) *i*, 𝑆 is a subset of the attributes (SNPs) 𝐹, 𝑓_𝑆∪{*i*}_ is the predicted value from the prediction model with an attribute subset including *i*, 𝑓_𝑠_ is the predicted value from the prediction model without the attribute *i*. Since the Shapley scores were also calculated element-wise, 30 individuals were randomly selected from the test sets and averaged throughout the prediction scenarios. Since the Shapley scores were calculated from possible combinations with other attributes, the scores can be decomposed at the pairwise level to calculate the pairwise Shapley scores to be utilised as genomic marker-by-marker interaction effects.

For the fully connected and data-driven prior knowledge GAT models, their estimated attention values from Equation (1) were extracted for comparison as inferred genomic marker-by-marker interaction effects. The comparison between the two GAT models highlights differences in the captured predictive patterns, enabling the analysis of the effects of data-driven prior knowledge on the GAT prediction mechanism.

### 5. Gene network inference

The extracted genomic marker effects from the GAT models were projected to their corresponding genomic marker regions, accompanied by the selected genomic marker-by-marker interaction effects from RF. These gene networks were visualised in the form of circos plots (Krzywinski et al., 2009) as the final phase in EasiGP (Tomura et al., 2025a). The mean value of the genomic marker effects was leveraged to infer the gene networks from the naïve ensemble-average model. The captured gene networks were compared against the key gene regulators identified in previous studies: QTL information from Chen et al. (2019) for the TeoNAM dataset, Buckler et al. (2009) for the MaizeNAM dataset and the maize adaptation study for flowering time by Wisser et al. (2019).

### 6. Genomic prediction model evaluation

The prediction accuracy of the genomic prediction models was measured by Pearson correlation, measuring the accordance level of ranking between the observed and predicted phenotype values. A Pearson correlation of 1 indicates that both the predicted and observed phenotype values are completely accordant and hence perfectly accurate. The prediction error was measured by mean squared error (MSE). An MSE of 0 indicates no difference between the observed and predicted values and thus no squared prediction error.

The Diversity Prediction Theorem (Page, 2018) was used to measure the level of diversity in predicted phenotypes from each GAT model in the naïve ensemble-average model:

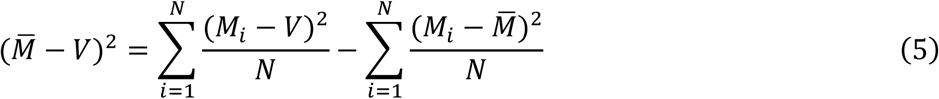

where *M*_*i*_ is the predicted value for the RILs from the prediction model (a GAT model) *i*, *M̅* is the mean predicted value from the *i* GAT models, *V* is the true value and *N* is the total number of individual prediction (GAT) models. In this study, observed phenotypes were used as *V*. The Many-Model ensemble error (first term) is calculated by subtracting the prediction diversity (third term) from the mean error of the individual prediction models (second term). Hence, Equation (5) shows that the ensemble Many-Model error decreases as the prediction values from the individual prediction models become more diverse.

The prediction performance of the genomic prediction models was iteratively evaluated under multiple prediction scenarios. For the TeoNAM, data were uniquely split into training and test sets 500 times per population and trait. Five training-test set ratios (0.8-0.2, 0.65-0.35, 0.5-0.5, 0.35-0.65 and 0.2-0.8) were used to investigate the effect of training set size. Hence, 12,500 prediction scenarios (5 populations * 5 ratios * 500 samplings) were generated per trait. For the MaizeNAM, data was sampled 50 times per population with the same five training-test set ratios. In total, 6,250 prediction scenarios (25 populations * 5 ratios * 50 samplings) were generated per trait.

## Results

### 1. Data-driven prior knowledge GAT models did not always improve overall prediction performance

Our results were not consistent with our hypothesis that the data-driven prior knowledge model would always outperform the other models across traits and datasets (Figure 2; Table S3). The prior knowledge model reached the highest median prediction accuracy for DTA in the MaizeNAM dataset, the lowest median prediction error for ASI in the MaizeNAM dataset and the lowest median prediction error for DTA in both datasets within the GAT models. However, its outperformance was not consistent in other prediction scenarios and metrics. The data-driven prior knowledge based on the Shapley scores generated for the RF graph models in this study did not always improve the prediction performance of the GAT models.

**Figure 2:**
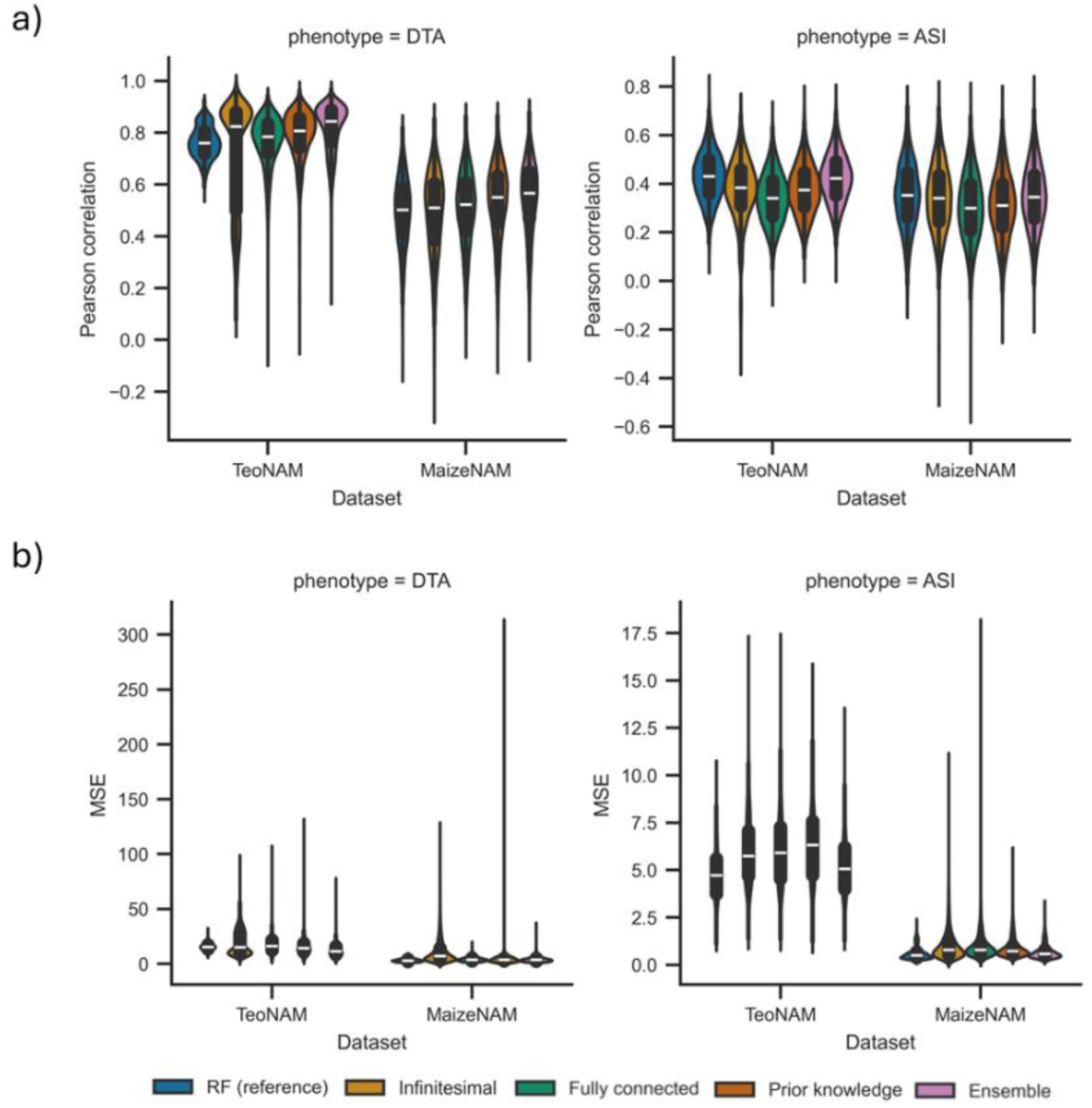
The comparison of prediction performance for infinitesimal, fully connected, data-driven prior knowledge and the naïve ensemble-average model (Ensemble) for days to anthesis (DTA) and anthesis to silking interval (ASI) traits in the TeoNAM and MaizeNAM datasets. Random forest was implemented as a reference (RF (reference)) used to infer the gene network as data-driven prior knowledge for the prior knowledge model. The prediction performance was measured in a) Pearson correlation for prediction correlation and b) mean squared error (MSE) for prediction error. For each trait, the prediction performance was measured in 12,500 prediction scenarios for the TeoNAM dataset and 6,250 prediction scenarios for the MaizeNAM dataset. The prediction scenarios were generated by the combination of the five training-test ratios (0.8- 0.2, 0.65-0.35, 0.5-0.5, 0.35-0.65 and 0.2-0.8), populations (5 and 25 for the TeoNAM and MaizeNAM datasets, respectively) and the total number of random dataset splits (500 and 50 for the TeoNAM and MaizeNAM datasets, respectively). The violin width represents the distribution of performance metrics. The white horizontal lines on the black box plots indicate the median value for each metric. The whiskers extend 1.5 times the interquartile range.

For DTA, the GAT model with the highest prediction performance was heavily dependent on the dataset (Figure 2; Table S3). In the TeoNAM dataset, the infinitesimal model reached the highest median prediction accuracy (Pearson correlation), while the prior knowledge model reached the lowest prediction error (MSE). In the MaizeNAM dataset, the prior knowledge model reached the highest median prediction accuracy and lowest prediction error. The prediction performance of the prior knowledge model was the highest or the second highest for most cases of DTA prediction within the GAT models.

The level of prediction performance improvement using data-driven prior knowledge was also dataset dependent for ASI (Figure 2; Table S3). In the TeoNAM dataset, the highest median prediction accuracy and lowest prediction error were achieved by the infinitesimal model within the GAT models. In the MaizeNAM dataset, this outperformance of the infinitesimal model was also observed in prediction accuracy, while the prior knowledge model reached the lowest prediction error within the GAT models. These prediction results indicate that the use of data-driven prior knowledge did not consistently improve the prediction performance of the GAT models for ASI.

### 2. Non-infinitesimal models showed greater robustness to limited training data

Prediction performance deterioration by reducing the training set size was less severe for the two non-infinitesimal GAT models (Figure 3). As an overall trend, the rate of reduction of the mean prediction performance increased as the size of the training set reduced for the infinitesimal model. For the TeoNAM dataset, such a trend was more strongly emphasised in DTA for both prediction accuracy and error, showing a greater performance reduction rate for the infinitesimal model after the training ratio became smaller than 0.5. In contrast, the other GAT models and their ensemble maintained the same reduction rate as the training set size became smaller. While the infinitesimal model maintained a stable prediction accuracy reduction rate by showing a similar reduction trend to the other models for ASI, the reduction rate became steeper after the training ratio of 0.35 for prediction error. For the MaizeNAM dataset, a similar trend of rapid performance decrease for the GAT infinitesimal model was observed as the training set size became smaller for both traits. The infinitesimal model showed an increased rate of performance reduction after the training ratio of 0.35 for DTA, measured with prediction error. The performance reduction rate was milder for ASI in prediction accuracy, as observed for the other GAT models. However, the prediction error increase of the infinitesimal model became more noticeable when the training size ratio was smaller than 0.5. Meanwhile, the other GAT models and the ensemble showed a more gradual increase in the performance reduction rate compared to the infinitesimal model. These results indicate that although the training set size generally reduced the prediction performance of all GAT models, explicitly specifying interactions between genomic markers reduced the decay of prediction performance when the training set size became smaller for DTA.

**Figure 3:**
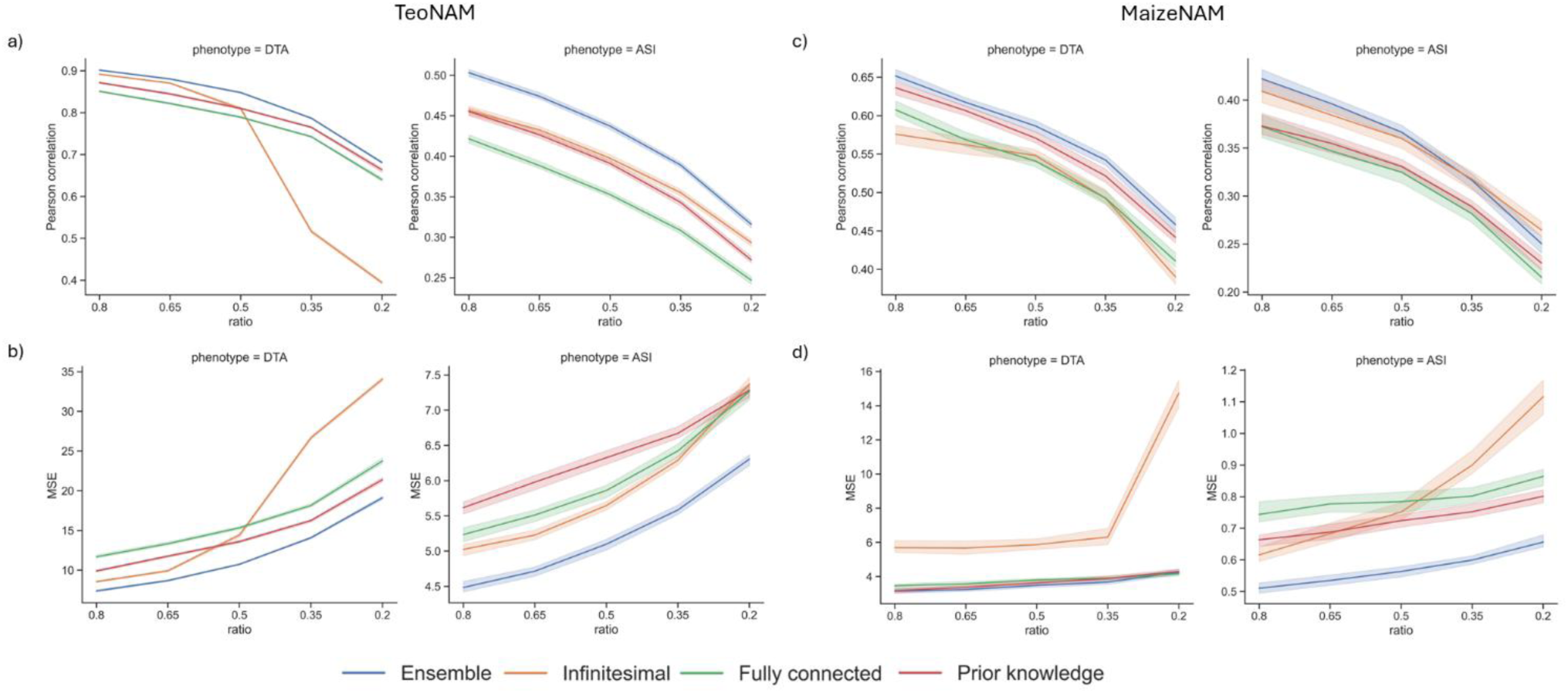
Prediction performance transition of the three GAT models (infinitesimal, fully connected and data-driven prior knowledge) and their ensemble as the ratio of the training set becomes smaller (0.8, 0.65, 0.5, 0,35 and 0.2), measured by a) Pearson correlation for the TeoNAM dataset, b) mean squared error (MSE) for the TeoNAM dataset, c) Pearson correlation for the MaizeNAM dataset and d) MSE for the MaizeNAM dataset. The upper and lower boundaries accompanying each line show the metric value range with standard error based on 2,500 and 1,250 prediction scenarios in each ratio for the TeoNAM and MaizeNAM datasets, respectively.

### 3. Ensembles of GAT models improved prediction performance

The ensembles of GAT models consistently outperformed or were comparable to the best GAT model (Figure 2; Table S3). In the TeoNAM dataset, the median prediction performance of the naïve ensemble-average model was higher than the GAT model with the highest prediction performance for both DTA and ASI. Similarly, the outperformance of the naïve ensemble-average model was observed for both DTA and ASI in the MaizeNAM. These prediction performance results indicate that the naïve ensemble-average model can consistently improve prediction performance relative to the individual GAT models.

The ensemble of the GAT models improved the prediction performance of the GAT-based approach, becoming comparable to RF used as a reference in this study. The prediction accuracy of the naïve ensemble-average model was also higher than that of RF in most cases. While the prediction error of RF was the highest in most cases, the difference from the ensemble was small compared to the individual GAT models. The result indicates that the ensemble of the GAT models in this study was an effective approach to improve prediction performance across the range of scenarios based on the datasets and traits.

### 4. Diversity of predictive information associated with increased prediction accuracy using ensembles

The increased diversity in the predicted phenotypes improved the prediction performance of the naïve ensemble-average model (Figure 4, Table S4). As the total number of GAT models included in the naïve ensemble average model increased, the mean ensemble error became lower and the mean Pearson correlation became higher in both DTA and ASI, especially for the TeoNAM dataset. Similarly, increasing the number of GAT models in the ensemble reduced the mean MSE. This strong association between the metrics indicates that the addition of the diverse GAT models into the naïve ensemble-average model tends to amplify the prediction performance of the naïve ensemble-average model.

**Figure 4:**
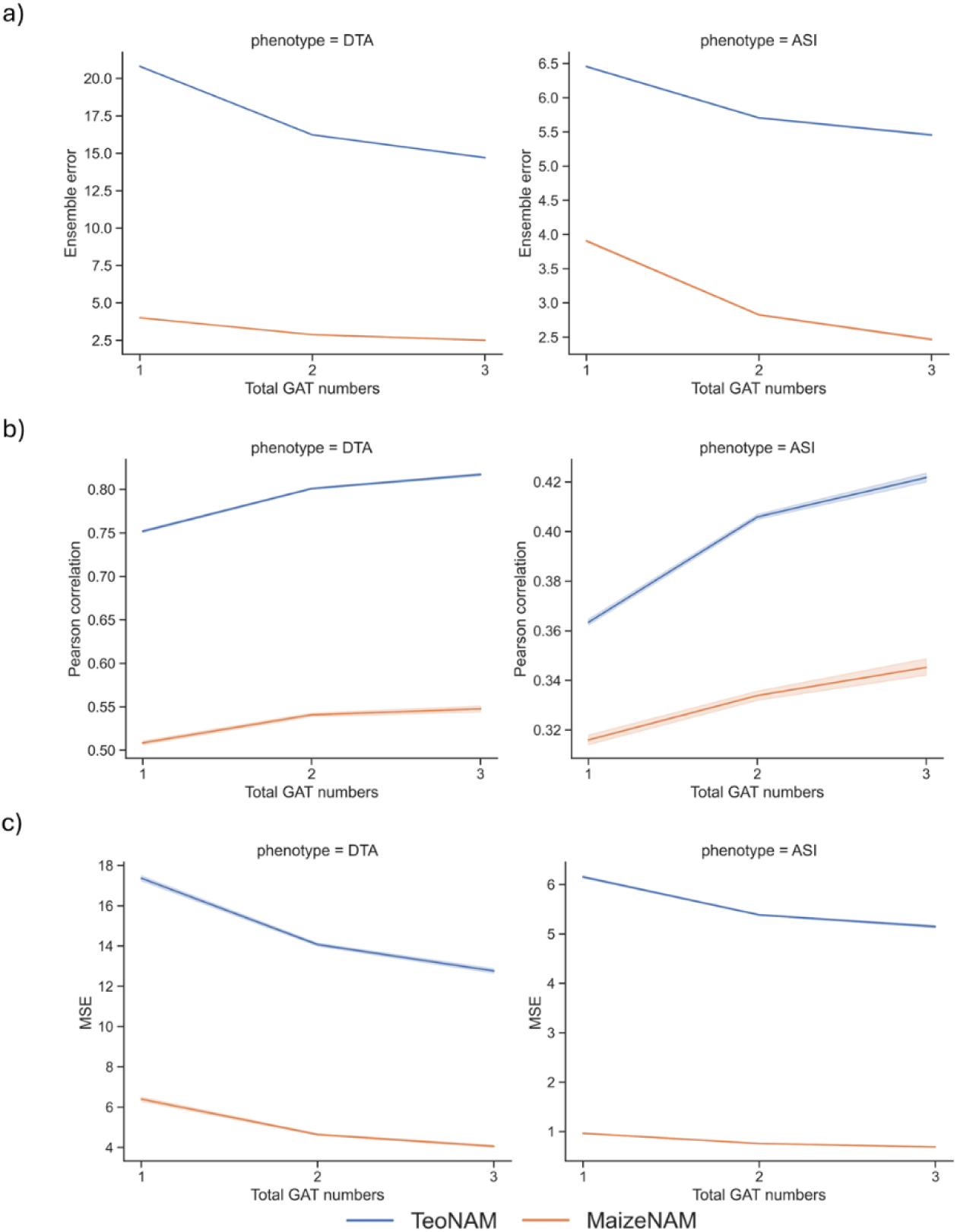
The metric transition of the naïve ensemble-average models with differences in the total number of included GAT models for the mean a) ensemble error in the Diversity Prediction Theorem, b) Pearson correlation and c) mean squared error (MSE) in both TeoNAM and MaizeNAM datasets. The total number of naïve ensemble-average models was 7, considering all the possible combinations of the GAT models that include single GAT models. Numbers 1, 2 and 3 of the x-axis represent the total number of the GAT models included in the naïve ensemble-average model. The upper and lower boundaries accompanying each line show the metric value range with standard error based on 12,500 for the TeoNAM dataset and 6,250 prediction scenarios for the MaizeNAM dataset.

The diversity in the captured information from each GAT model was also observed in the prediction outcome of the GAT models at both the phenotype and genome levels (Figure 5, S1; Table S5). For DTA at the phenotype level, the mean correlation values between GAT pairs ranged from 0.66 to 0.85 for the TeoNAM dataset, while the range was from 0.79 to 0.93 for the MaizeNAM dataset. The lower correlation indicates higher diversity in predicted phenotype values between GAT models. For ASI at the phenotype level, the mean correlation values ranged from 0.59 to 0.67 for the TeoNAM dataset, while the range was from 0.56 to 0.76 for the MaizeNAM dataset. Following the implication of the Diversity Prediction Theorem, lower correlation (high diversity) at the phenotype level contributes to higher prediction performance improvement by ensembles. These observed correlation values indicated that diversity existed at the phenotype level, contributing to performance improvement using the naïve ensemble-average model. For the genome level, diversity was emphasised for both traits and datasets. Diversely captured genome-level information contributed to diverse predicted phenotypes across the prediction scenarios. The diversity of the GAT models at the genome level was also observed in the generated circos plots (Figure 6, S2). The patterns of the colour intensity in each ring corresponding to the magnitude of genomic marker effects varied for each GAT model. Variation in the marker effects indicates that the GAT models captured features of the standing genetic variation differently and the ensemble generated a unique view of the gene network by integrating the distinctively estimated additive and non-additive genomic marker effects included in the standing variation from the GAT models.

**Figure 5:**
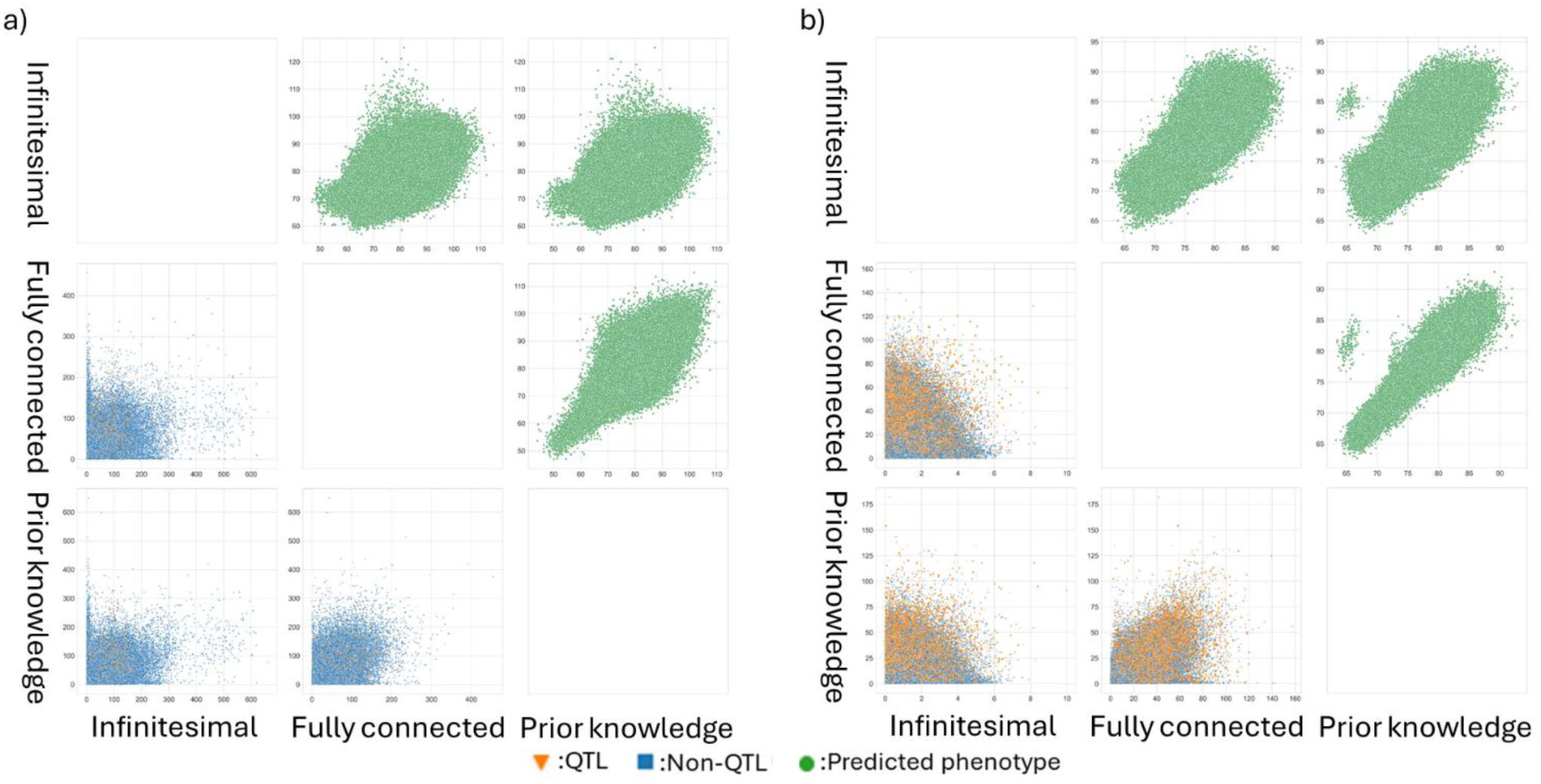
Pairwise comparisons of the GAT models (infinitesimal, fully connected and data-driven prior knowledge) for the days to anthesis (DTA) trait across all the prediction scenarios for a) the TeoNAM (12,500 prediction scenarios) and b) MaizeNAM (6,250 prediction scenarios) datasets at the phenotype and genome levels. Within each subplot in the scatter plot matrix, the GAT models were compared for predicted phenotypes (top right triangle) and genomic marker effects (the bottom left triangle). The green dots represent a pair of predicted phenotypes for RIL samples in the test sets for each prediction scenario. The blue squares and orange triangles represent pairs of inferred effects of genomic markers in each sample scenario for non-QTL and QTL, respectively. Non-QTL and QTL markers were identified by Chen et al. (2019) for the TeoNAM dataset and Buckler et al. (2009) for the MaizeNAM dataset.

**Figure 6:**
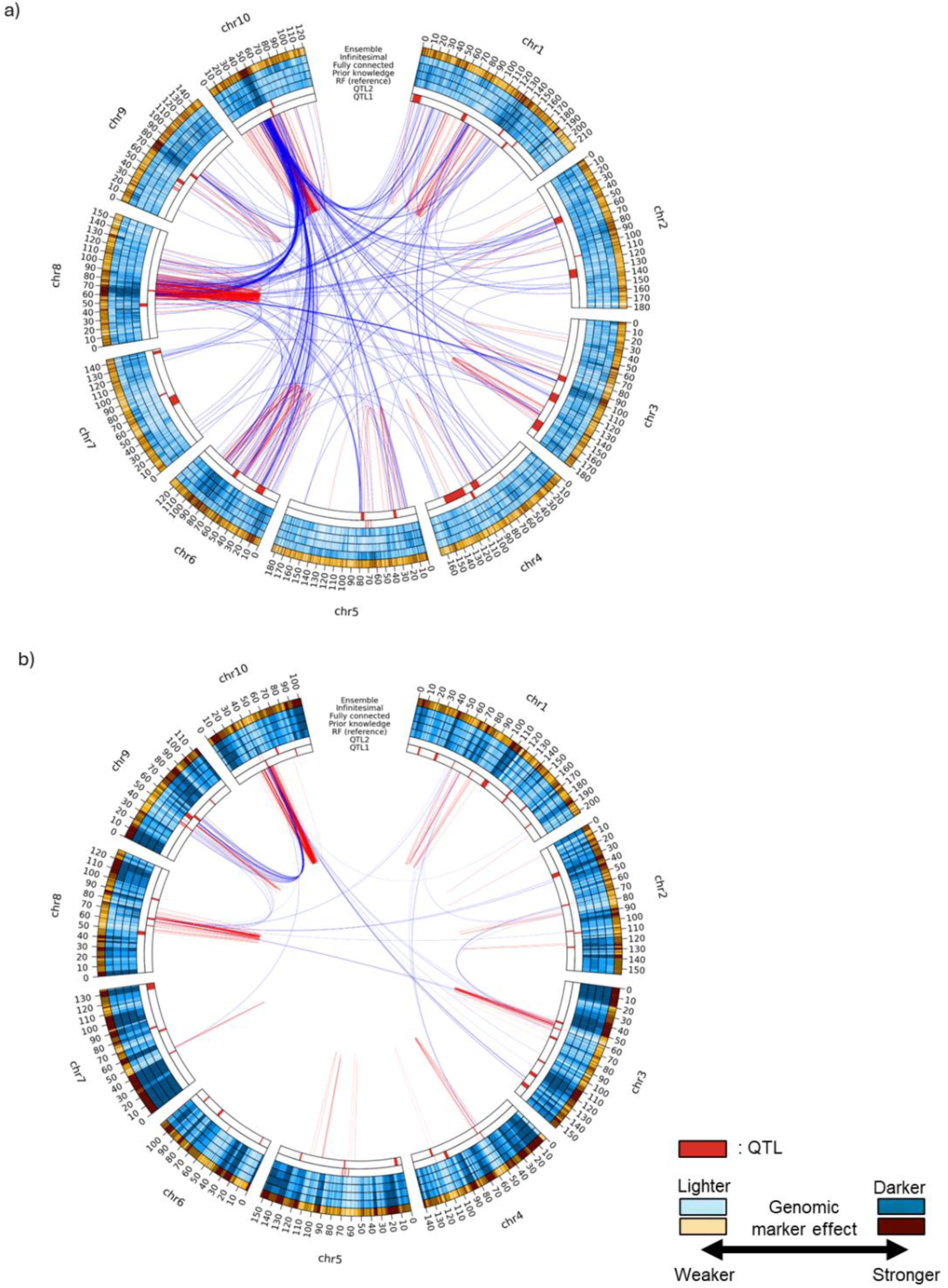
Circos plots for the days to anthesis (DTA) trait for the a) TeoNAM and b) MaizeNAM datasets. The innermost ring (QTL1) represents QTL regions in red identified by Chen et al. (2019) for the TeoNAM dataset and Buckler et al. (2009) for the MaizeNAM dataset, respectively. The second innermost ring (QTL2) represents the QTL regions identified by Wisser et al. (2019). The subsequent four blue rings represent the genomic marker effects from random forest as reference (RF (reference)) used to extract data-driven prior knowledge, the data-driven prior knowledge model, the fully connected model and the infinitesimal model. The outermost ring in orange shows the estimated genomic marker effects of the naïve ensemble-average model. The intensity of the colours shows the strength of the genomic marker effects represented in ten levels based on the quantiles. The colour becomes darker as the genomic marker effects become higher. Links between two genomic marker regions represent genomic marker-by-marker interactions detected by RF using the Shapley scores. The genomic marker pairs with the Shapley scores included in the top 0.001% were selected for the TeoNAM dataset and 0.05% for the MaizeNAM dataset, considering the difference in the total number of identified marker combinations. Thicker lines show stronger marker-by-marker interactions. The red and blue lines represent genomic marker-by-marker interactions within and between chromosomes, respectively. The genetic distance is represented in centimorgans (cM).

Such diversity in the captured standing genetic variation was also represented in the distribution of attention values from the non-infinitesimal GAT models (Figure 7). Comparison of the attention value histograms between the GAT fully connected and data-driven prior knowledge models showed that the prior knowledge model assigned larger attention values to specific edges, represented as a more right-skewed distribution. In particular, this tendency was emphasised for DTA in the MaizeNAM dataset, showing a right-skewed distribution for the prior knowledge model, while a normal distribution was observed for the fully connected model. This result indicates that the data-driven prior knowledge model was likely to learn several key predictive patterns by highlighting specific edges, especially for the MaizeNAM dataset, contributing to the higher overall prediction performance of the prior knowledge model for the MaizeNAM dataset compared to the other GAT models. Since the non-infinitesimal models differently highlighted key edges connecting genomic marker nodes, the identified key predictive interaction patterns were diverse (Figure S3, S4). Such diverse interaction patterns also consequently contributed to improved prediction performance by the ensemble.

**Figure 7:**
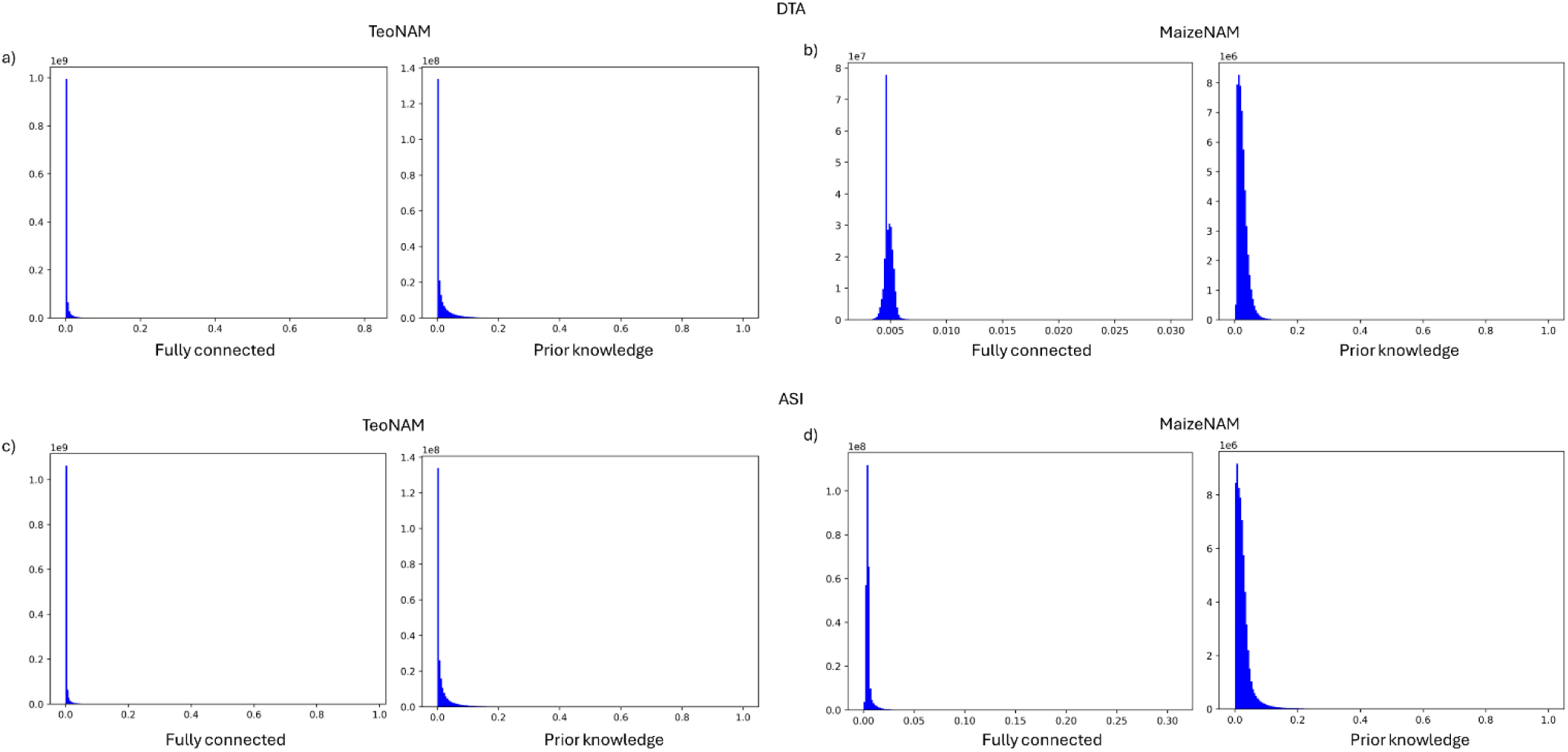
The comparison of attention values extracted from the GAT fully connected and GAT data-driven prior knowledge models. The histograms represent the distribution of the attention values for a) the days to anthesis (DTA) for the TeoNAM dataset, b) DTA for the MaizeNAM dataset, c) the anthesis to silking interval (ASI) for the TeoNAM dataset and d) ASI for the MaizeNAM dataset.

### 5. GAT models identified several key gene regulators

Highlighted genomic marker regions from the GAT models overlapped with several key gene regulators for DTA that had been identified in previous studies (Figure 5). For instance, the genomic marker region between 35 cM and 50 cM in chromosome 10 was identified across genomic prediction models for DTA in the TeoNAM dataset. This region is in close proximity to *ZmCCT10* (55 cM), known to be one of the most significant maize flowering time genes through its control of the photoperiod pathway (Dong et al., 2012; Chen et al., 2019; Wisser et al., 2019). This highlighted region in chromosome 10 showed heavy interactions with another region containing the gene *ZCN8* (70 cM in chromosome 8), captured by all the GAT models. *ZCN8* is another gene that integrates signals from the photoperiod pathway, regulating the transition process from the vegetative to reproductive stage (Meng et al., 2011; Dong et al., 2012; Guo et al., 2018; Aiyesa et al., 2025). For the MaizeNAM dataset, the highlighted genomic marker region overlapped with the gene region homologue of *Ghd7* (85 cM in chromosome 1) in rice (Xue et al., 2008; Buckler et al., 2009). The GAT models also highlighted the genomic marker region close to *ZCN8*, as observed in the TeoNAM dataset, and further recognised the marker region close to another photoperiod pathway gene, *ZmCCT9* (55cM in chromosome 9). The photoperiod pathway gene *ZmCCT9* regulates flowering time through responses to the changes in daylength associated with levels of latitude by controlling the expression of *ZCN8* (Huang et al., 2018). Such overlaps were frequently observed with known gene regulators reported in the literature aligning with markers assigned large effects in the GAT genomic prediction models for both DTA and ASI traits (Figure 5, S2), indicating that the genomic prediction models included genomic regions containing genes with an established influence on target traits for their prediction.

## Discussion

### 1. More comprehensive prior knowledge may improve the prediction performance of GAT models

The primary objective of this study was to investigate the possibility of performance improvement in genomic prediction through the integration of data-driven gene networks as prior knowledge over other G2P structures, analysed through the implementation of GAT models. A lack of consistency in prediction performance improvement has remained a critical issue despite the introduction of numerous prediction algorithms, attributed to the effect of the No Free Lunch Theorem (Wolpert and Macready, 1997). The integration of data-driven prior knowledge was expected to mitigate this problem by customising genomic prediction models into problem-specific ones.

Contrary to the expected outcome, there was still a lack of consistency in the overall prediction performance improvement of the GAT models using data-driven prior knowledge (Figure 2; Table S3). There are a number of possible reasons. This inconsistent performance improvement using data-driven prior knowledge might have been attributed to the potential imprecision in the captured genomic marker-by-marker interactions. Shapley scores were estimated using SHAP (Lundberg and Lee, 2017) incorporated in EasiGP, approximating Shapley values as “scores” instead of calculating the actual values to overcome the prohibitive computational time by leveraging conditional expectations and a perturbation approach (Janzing et al., 2020; Molnar et al., 2020; Contreras et al., 2024). A series of approximation formulas, such as kernel-based approaches, allows the estimation of Shapley values in a computationally efficient manner (Lundberg and Lee, 2017). However, such an approximation approach may calculate Shapley scores that may not be aligned with actual Shapley values, especially when a number of irrelevant attributes were included in the datasets (Huang and Marques-Silva, 2024). The approximation formulas can be affected by the irrelevant attributes that can distort Shapley scores as noise. The leveraged perturbation approach may also affect this imprecision by considering the attribute value combinations that do not reflect real-world conditions (Contreras et al., 2024). Hence, Shapley scores may not represent the actual influential size of each attribute (genomic marker). Inaccurate Shapley scores may have compromised the effectiveness of the supervision of the GAT models in this study.

To capture more informative prior knowledge from data, genomic marker-by-marker interactions can be inferred as epistasis using conditional dependencies between the markers (Behrouzi and Wit, 2019). This approach only extracts genomic marker pairs that are directly conditionally dependent, removing noncausal relationships created by any genomic marker pair that is conditionally independent but correlated through another genomic marker that is highly correlated with the two. This alternative approach may help capture more precise relationships among genomic markers. Exclusion of noncausal connections was not explicitly implemented in the current Shapley score-based method. Another approach for estimating genomic marker-by-marker interactions is BShap (Sundararajan and Najmi, 2020), which extends Shapley scores using a baseline. The baseline returns a prediction value excluding the effect of a target attribute (genomic marker). The contribution of the target attribute is calculated as the difference between the prediction values including and excluding the target attribute, as employed in Integrated Gradients. These potential approaches may vary in how they weigh genomic marker-by-marker interaction effects, consequently generating diverse views of the standing genetic variation. Thus, integration of all these approaches into an ensemble might enhance prediction performance by providing more comprehensive prior knowledge inferred from data to assist the GAT models. Alternative approaches for marker-by-marker interaction effect inference are an area for further investigation as indicated by this study.

The complexity of genomic marker-by-marker interactions might also have affected the informativeness of the data-driven prior knowledge. ASI is a secondary trait (Leng et al., 2022) calculated by the difference between DTA and DTS and thus can be regulated by the gene network of both quantitative traits with other crop development traits (Messina et al., 2019). Consequently, ASI might be influenced by the combination of two or more distinct gene networks, thereby increasing the complexity of the trait gene network. The current inferred gene network likely did not capture such complex interactions between genomic markers in ASI due to the lower prediction performance of RF used to calculate the Shapley scores. Hence, the integrated data-driven prior knowledge may not have improved the prediction performance of GAT for ASI regulated by potentially more complex gene networks.

The extension of prior knowledge using biological prior information potentially captures such biological complexity regulating the target traits. For instance, omics data from transcriptomics, metabolomics and proteomics can be integrated into genomic data layers. Each component of omics data can be represented as a graph based on empirical verification and statistical analysis, combined with others to develop an extensive biological graph. The effect of the comprehensive level of the biological graph on GAT prediction performance can be evaluated by sequentially adding components of the omics graphical data within the hierarchical G2P model, as discussed by Powell et al. (2022). Such a comparative study helps build an understanding of the association between the comprehensive level of biological prior knowledge and the improvement level of GAT prediction performance. This analysis determines the optimal level of biological information for target trait prediction, considering the cost from a practical view in crop breeding programs. Further, the performance of any GAT model incorporating additional layers of omics data can be compared to the developed baseline GAT model applying the methods demonstrated herein. For genomic level data, the data-driven gene networks in this study can be extended by combining the well-known key gene interactions. While the current data-driven approach focused on a data-driven definition of a prior graph topology by applying an RF approach to extract the overall interactions between genomic markers, some specific critical gene interactions may not be detected by this approach. In contrast, only using the empirically identified gene networks might fail to capture unknown interactions. Hence, reliable gene networks can be combined with the current data-driven gene networks to potentially improve the prediction performance of GAT prior knowledge models (Marjoram et al., 2014; von Rueden et al., 2023; Messina et al., 2025). To date the potential of such biology-based prior knowledge approaches has been investigated mainly in human medicine (Xiao et al., 2023; Alharbi et al., 2025) but has not yet been deeply researched for applications to enhance crop genomic prediction. Using the GAT methodology demonstrated in this study as the foundation, there are new opportunities to investigate the effects of incorporating biological prior knowledge on prediction performance. These will be targets for future investigations as suitable datasets become available.

### 2. An ensemble of GAT models can consistently improve prediction performance

While the methods discussed above to improve the performance of the GAT models using data-driven prior knowledge, based on the Diversity Prediction Theorem expectations, they may still not outperform an ensemble-based approach. The prediction performance of the ensemble was consistently higher than or equivalent to the best GAT models and the RF baseline (Figure 2, S3). This consistent prediction performance increase by the naïve ensemble-average can contribute to improving prediction performance across many prediction scenarios in crop breeding programs. Such an approach is especially useful given the difficulty in selecting the optimal genomic prediction model when prediction performance is heavily scenario-dependent (Voss-Fels et al., 2019; Escamilla et al., 2025). Consequently, it has been a challenge for breeders to achieve high prediction performance across all of their target trait by dataset prediction scenarios. Our results suggest that the risk of selecting an unsuitable genomic prediction model can be mitigated by an ensemble of GAT models, which consistently match or outperform the individual models.

Such performance improvement by the ensemble was likely attributed to diversity in phenotypic predictions (Figure 5, S1) as elucidated in the Diversity Prediction Theorem (Page, 2018; Messina et al., 2025). When diversity increased by adding more GAT models to the ensemble, the prediction performance of the ensemble improved, with increases in prediction accuracy and reductions in error (Figure 4, Table S4). This positive association between diversity and prediction performance of the ensemble was also observed in other studies (Dietterich, 2000a; Melville et al., 2005; Kick and Washburn, 2023), showing a downward trend in prediction error as the number of individual prediction models included in the ensemble increased. The aggregation of diverse prediction models can offset prediction errors and weaknesses of each, improving the prediction performance of the ensemble approach (Kick and Washburn, 2023; Washburn et al., 2025). The diversity in the ensemble might support reaching the global optimum rather than convergence at the suboptimal local optimum (Bourel et al., 2024). Complex prediction models, including GAT, aim to solve optimisation problems (reaching the highest possible prediction performance) using a training set in a non-convex way that contains numerous local optima. There is a risk that the prediction performance of such complex prediction models can be optimised locally, and the model parameters may not be further adjusted for performance improvement (Audhkhasi et al., 2012). GAT may stop a drastic adjustment of trainable weights once it starts reaching a convergence in the prediction performance improvement, but the global optima can be achieved by another set of weights that is far from the current local optima. This problem can also be described as overfitting, insufficiently generalising the predictive power (Karystinos et al., 2000). The stagnation at the local optima can be mitigated by combining diverse local optima through the ensemble approach (Dietterich, 2000b). Hence, the ensemble of the multiple GAT models improved prediction performance by approaching the global optima.

The performance improvement using diverse predictive information in the ensemble was also demonstrated at the genome level by each GAT model assigning a different effect contribution for each genomic marker (Figure 5, 6, 7, S1, S2, S3, S4). By combining the distinct inferred gene networks from each GAT model through the ensemble, a more comprehensive fraction of the standing trait genetic variation could be captured to enhance prediction performance (Cooper et al., 2025; Messina et al., 2025; Tomura et al., 2025b). The interpretable genomic prediction models enabled such investigation at the genome level in this study, which has not been well-investigated in the realm of genomic prediction for crop breeding. The significance of accurately capturing the standing variation was aligned with the conclusion of Morgante et al. (2018), showing that there was a prediction performance improvement when the true standing variation of trait genetic architecture was well-captured in simulated data. In contrast, prediction performance remained low when the true standing variation of the trait genetic architecture was not clearly identified and heavy interactions between genomic markers (fully connected-like structure) were included. Hence, capturing a gene network containing a comprehensive view of standing variation that closely reflects the true genetic architecture is a key step in improving the predictive performance of genomic prediction models (Cooper et al., 2005).

The use of simulated datasets may complement the empirical investigations and help understand the potential of the prediction performance of GAT in relation to the properties of trait gene networks as future research. The true trait genetic network underlying data can be predefined in the simulation datasets, enabling the development of a series of simulation datasets that gradually increase the complexity level of the trait gene network (Cooper et al., 2005). Simulation data can also be further extended to the omics levels. For instance, Powell et al. (2022) simulated the trajectory of changes in the time to bud outgrowth grounded in the biological knowledge from the interactions between hormones and sucrose, with the effects of simulated causal genetic loci. The continuum of complexity level in the generated trait gene and omics networks facilitates the association analysis between the precision of the detected trait biological network by GAT models and their prediction performance.

### 3. Genomic marker-by-marker interactions can supplement information loss for small training sets

The prediction performance of the infinitesimal model was more reactive to the size of the training set. The reduction rate of prediction performance was larger than that of the fully connected or data-driven prior knowledge models when the size of the training set became smaller, especially for DTA in the TeoNAM dataset and ASI in the MaizeNAM dataset (Figure 3). Maintaining prediction performance in the non-infinitesimal models might have been due to the simplicity of the provided graph structures in the GAT infinitesimal model. It might have been relatively easy for the infinitesimal model to learn key prediction patterns with a large training set due to the absence of explicit complex interactions between genomic markers in their graph structures. Consequently, the infinitesimal model outperformed the other models. However, the simplicity in the graph structure reduced the effectiveness of the infinitesimal models in learning key predictive patterns when the allocated training set was small. There is a tendency that a star-shaped (infinitesimal) graphical structure can cause overfitting as the training set becomes smaller (Bechler-Speicher et al., 2024). In contrast, the explicit marker-by-marker interactions might have supplemented the information loss derived from the diminishing size of the training set for the non-infinitesimal models. These explicit marker-by-marker interactions might have improved the prediction performance of the non-infinitesimal model when the training set size became smaller.

Among the non-infinitesimal models in this study, the integration of data-driven prior knowledge showed higher prediction performance as an overall trend. Unnecessary explicit interactions between genomic markers can introduce noise, thereby negatively affecting prediction performance. Selecting genomic marker-by-marker interactions using RF with Shapley scores might have excluded such unnecessary explicit interactions. Hence, prior knowledge can provide more accurate marker-by-marker interaction information into graphs, contributing to performance improvement especially when the training set is small (Schapire et al., 2002). In crop breeding programs, it can be a challenge to conduct large experiments based on a large number of individuals with their genotypic and phenotypic information recorded (Ibba et al., 2020; Sneller et al., 2021; Jines et al., 2025). Therefore, using prior knowledge information for the GAT models can be a preferable approach compared to the fully connected GAT models.

The use of inferred marker effects as prior knowledge is conceptually similar to the prediction mechanism of the standard ridge regression (Meuwissen et al., 2001) with a high penalty and Bayesian regression (Meuwissen et al., 2001) with a strong prior around 0 for many marker effects. However, a critical advantage of the prior knowledge model is the direct integration of topological information into the prediction mechanisms, given that the amount of training data is sufficiently large. GAT prior knowledge models can utilise the inferred topological information to effectively extract key predictive patterns as embeddings. Such topological information cannot be utilised by the conventional approaches trained on tabular data.

### 4. Interpretability of genomic prediction models can accelerate genetic gain

The interpretability of the GAT models and the ensemble approach can identify genomic marker regions that are highly influential on target traits. Identifying preferred allele combinations in the highlighted genomic marker regions using the interpretable GAT models helps breeders select individuals with desirable traits. The accordance rate of their alleles in those genomic marker regions can be used as a selection criterion, in contrast to considering all alleles across the genome. This potential selection criterion can be used simultaneously with other criteria, such as the ranking from predicted phenotypes and genetic diversity. The combination of multiple selection criteria can improve the selection performance of individuals with more desirable traits. Pook et al. (2025) accelerated the genetic gain of simulated maize breeding programs by developing a formula that combines several selection criteria from the view of short-term, long-term genetic gain and the level of genetic diversity. Hence, the combination of diverse selection criteria can also positively affect the speed of genetic gain.

The interpretable genomic prediction models also identified several new genomic marker regions. This interpretability-oriented genomic prediction approach has been applied to infer the gene network in other fields such as detecting genes influencing body heights and schizophrenia in humans (Askland et al., 2021; van Hilten et al., 2021; Hequet et al., 2025), muscle growth in livestock (Guo et al., 2024) and fungus tolerance in ash trees (Doonan et al., 2025). Interpretable machine learning models have started being used in crop breeding with the application of interpretable RF and deep learning models (Novielli et al., 2024; Wang et al., 2025b), but no other studies have collectively utilised interpretable genomic prediction models as an ensemble. Although validation of new genomic marker regions identified through the ensemble approach is still required, this strategy has the potential to help breeders focus on the identified gene regions rather than the comprehensive investigation of the whole genome to reduce cost and save time. It is encouraging that multiple regions containing previously identified key genes involved in controlling flowering time in maize were consistently identified across two divergent datasets, the TeoNAM and MaizeNAM, in the applications of GAT and their ensembles.

## Conclusion

The potential for developing GAT genomic prediction methodology for crop breeding applications has been empirically demonstrated. In this study, the prediction performance was evaluated for three GAT models incorporating different G2P structures relevant to genomic prediction for crop breeding, and ensembles of GAT models. The prediction results from our study showed that while using the GAT models with data-driven prior knowledge did not always increase the prediction performance, the ensemble of multiple GAT models, including the one with prior knowledge, consistently improved prediction performance. Capturing a more complete view of the standing genetic variation using the ensemble was likely a critical component for prediction performance improvement in the experimental scenarios. GAT models can be further investigated by integrating additional biological prior knowledge and simulation data for crop genomic prediction using the GAT methodology demonstrated in this study. Such extensions can potentially improve further the genomic prediction performance of GAT and thereby the ensemble, accelerating genetic gain by enhancing selection accuracy.

Data and code availability

The original TeoNAM dataset (Chen et al., 2019) used in this study is located at https://datacommons.cyverse.org/bro-wse/iplant/home/shared/panzea/genotypes/GBS/TeosinteNAM for the genotypes and https://gsajournals.figs-hare.com/articles/dataset/Supplemental_Material_for_Chen_et_-al_2019/9250682 for the phenotypes. The original MaizeNAM dataset (Buckler et al., 2009) used in this study is located at https://cbsusrv04.tc.cornell.edu/use-rs/panzea/download.aspx?filegroupid=10 for the genotypes and phenotypes. The computational tool, EasiGP, and the downloaded datasets are located at https://github.com/ShunichiroT/EasiGP.

## Acknowledgments

We thank the National Computational Infrastructure (NCI) and the Research Computing Centre (RCC) at the University of Queensland for providing access to the High Performance Computing (HPC) machines. We would also like to thank the editor and two anonymous reviewers for their excellent suggestions to improve the manuscript.

## Conflicts of interest

The authors declare no conflicts of interest.

Supplementary Material

## Supplementary Material

**Table S1:** The number of genomic markers (SNPs) at the original and after linkage disequilibrium (LD) filtering and records from recombinant inbred lines (RILs) in the TeoNAM and MaizeNAM datasets. Each subpopulation contains a different total number of SNPs in the TeoNAM dataset. The total number of SNPs after LD filtering depends on the combination of records included in the training set in each prediction scenario. The total number of records varies depending on the subpopulation for both datasets.

| Dataset | SNPs |  | Total records |
| --- | --- | --- | --- |
|  | Original | LD filtering |  |
| TeoNAM | 11,396 - 16,110 | 260 - 355 | 438 - 616 |
| MaizeNAM | 1,106 | 170 - 251 | 126 - 196 |

**Table S2:**
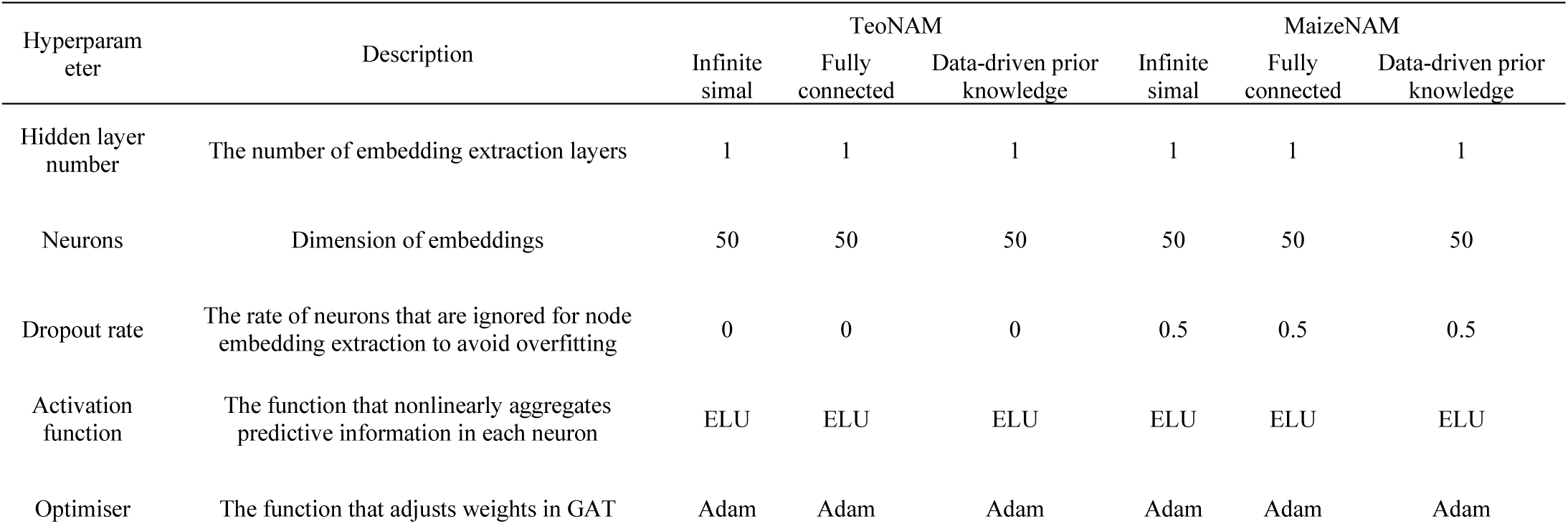
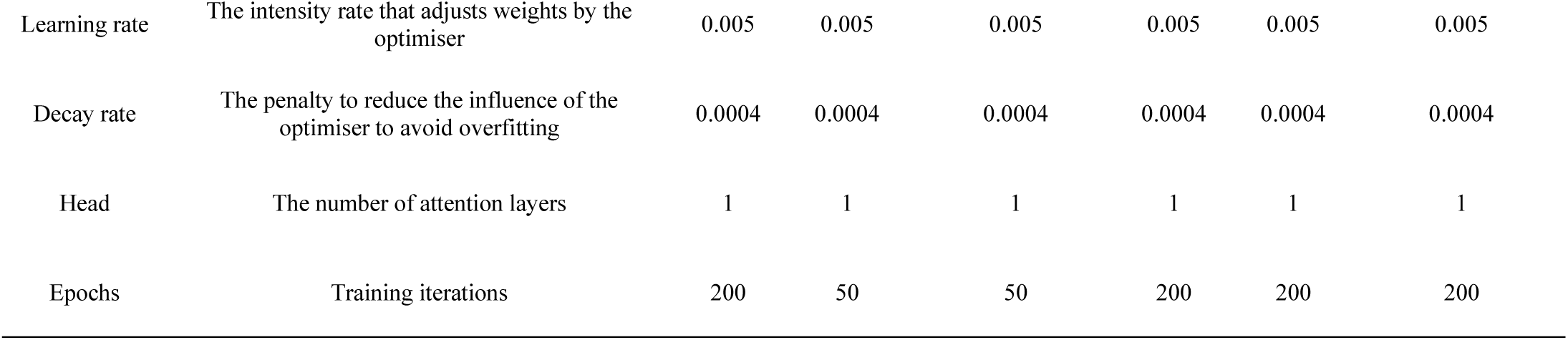
The selected hyperparameters for the GAT models in this study.

| Hyperparameter | Description | TeoNAM |  |  | MaizeNAM |  |  |
| --- | --- | --- | --- | --- | --- | --- | --- |
|  |  | Infinite simal | Fully connected | Data-driven prior knowledge | Infinite simal | Fully connected | Data-driven prior knowledge |
| Hidden layer number | The number of embedding extraction layers | 1 | 1 | 1 | 1 | 1 | 1 |
| Neurons | Dimension of embeddings | 50 | 50 | 50 | 50 | 50 | 50 |
| Dropout rate | The rate of neurons that are ignored for node embedding extraction to avoid overfitting | 0 | 0 | 0 | 0.5 | 0.5 | 0.5 |
| Activation function | The function that nonlinearly aggregates predictive information in each neuron | ELU | ELU | ELU | ELU | ELU | ELU |
| Optimiser | The function that adjusts weights in GAT | Adam | Adam | Adam | Adam | Adam | Adam |
| Learning rate | The intensity rate that adjusts weights by the optimiser | 0.005 | 0.005 | 0.005 | 0.005 | 0.005 | 0.005 |
| Decay rate | The penalty to reduce the influence of the optimiser to avoid overfitting | 0.0004 | 0.0004 | 0.0004 | 0.0004 | 0.0004 | 0.0004 |
| Head | The number of attention layers | 1 | 1 | 1 | 1 | 1 | 1 |
| Epochs | Training iterations | 200 | 50 | 50 | 200 | 200 | 200 |

**Table S3:**
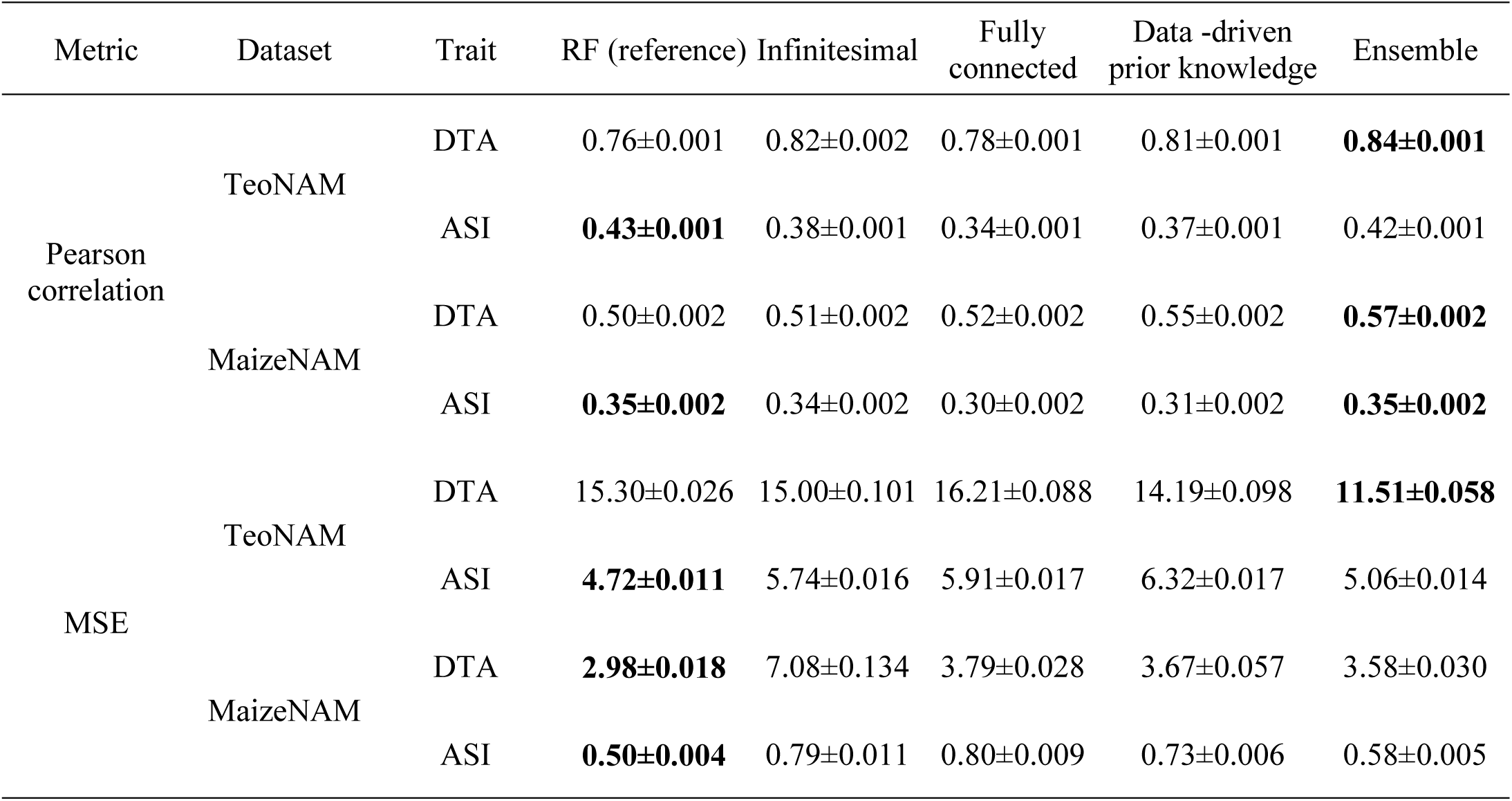
The prediction performance of each GAT model and RF as reference, measured in the median prediction accuracy (Pearson correlation) and median prediction error (mean squared error (MSE)) for the days to anthesis (DTA) and anthesis to silking interval (ASI) in the TeoNAM and MaizeNAM datasets. The sign “±” indicates the standard error of each median metric value. The highlighted values in black indicate the highest prediction performance in each scenario. The displayed median metric values correspond to Figure 2.

| Metric | Dataset | Trait | RF (reference) | Infinitesimal | Fully connected | Data -driven prior knowledge | Ensemble |
| --- | --- | --- | --- | --- | --- | --- | --- |
| Pearson correlation | TeoNAM | DTA | 0.76±0.001 | 0.82±0.002 | 0.78±0.001 | 0.81±0.001 | **0.84±0.001** |
|  |  | ASI | **0.43±0.001** | 0.38±0.001 | 0.34±0.001 | 0.37±0.001 | 0.42±0.001 |
|  | MaizeNAM | DTA | 0.50±0.002 | 0.51±0.002 | 0.52±0.002 | 0.55±0.002 | **0.57±0.002** |
|  |  | ASI | **0.35±0.002** | 0.34±0.002 | 0.30±0.002 | 0.31±0.002 | **0.35±0.002** |
| MSE | TeoNAM | DTA | 15.30±0.026 | 15.00±0.101 | 16.21±0.088 | 14.19±0.098 | **11.51±0.058** |
|  |  | ASI | **4.72±0.011** | 5.74±0.016 | 5.91±0.017 | 6.32±0.017 | 5.06±0.014 |
|  | MaizeNAM | DTA | **2.98±0.018** | 7.08±0.134 | 3.79±0.028 | 3.67±0.057 | 3.58±0.030 |
|  |  | ASI | **0.50±0.004** | 0.79±0.011 | 0.80±0.009 | 0.73±0.006 | 0.58±0.005 |

**Table S4:**
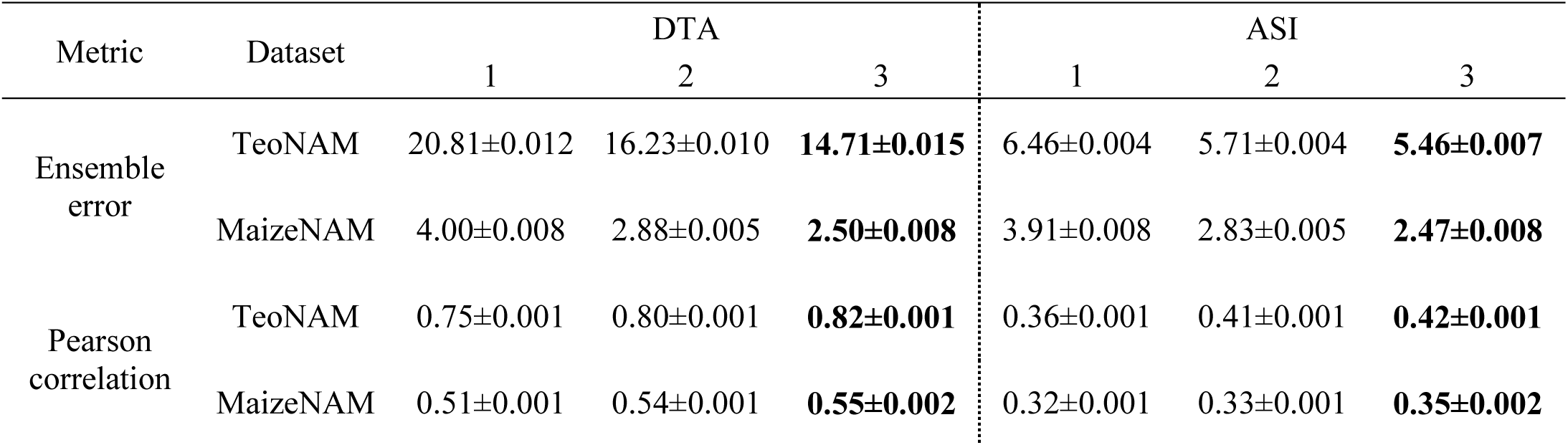
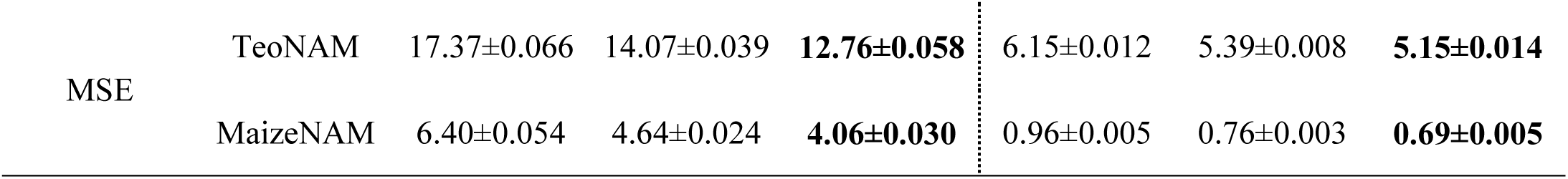
The evaluation of the naïve ensemble-average models with differences in the total number of included GAT models for the days to anthesis (DTA) and anthesis to silking interval (ASI) traits in both TeoNAM and MaizeNAM datasets, measured in the mean ensemble error from the Diversity Prediction Theorem, Pearson correlation and mean squared error (MSE). The total number of naïve ensemble-average models was 7, considering all the possible combinations of the GAT models that include single GAT models. Numbers 1, 2 and 3 of the columns represent the number of the GAT models included in the naïve ensemble-average model. The sign “±” indicates the standard error corresponding to each metric value. The highlighted values in black indicate the highest performance in each scenario. The displayed median metric values correspond to Figure 4.

**Table S5:** Pearson correlation of each GAT model (infinitesimal, fully connected and data-driven prior knowledge) pair at the level of the predicted phenotypes (the top right triangle) and the genomic marker effects (the bottom left triangle) in a) the days to anthesis (DTA) for the TeoNAM dataset, b) DTA for the MaizeNAM dataset, c) the anthesis to silking interval (ASI) for the TeoNAM and d) ASI for the MaizeNAM dataset. For a) and b), each correlation value in the table matrix corresponds to Figure 5, while for c) and d), each correlation value in the table matrix corresponds to Figure S1.

|  |  |  |  |  |
| --- | --- | --- | --- | --- |
| a) |  | Infinitesimal | Fully connected | Prior knowledge |
|  | Infinitesimal | - | 0.66 | 0.67 |
|  | Fully connected | 0.30 | - | 0.85 |
|  | Prior knowledge | 0.33 | 0.62 | - |
| b) |  | Infinitesimal | Fully connected | Prior knowledge |
|  | Infinitesimal | - | 0.79 | 0.78 |
|  | Fully connected | 0.05 | - | 0.93 |
|  | Prior knowledge | 0.07 | 0.68 | - |
| c) |  | Infinitesimal | Fully connected | Prior knowledge |
|  | Infinitesimal | - | 0.59 | 0.63 |
|  | Fully connected | 0.15 | - | 0.67 |
|  | Prior knowledge | 0.19 | 0.23 | - |
| d) |  | Infinitesimal | Fully connected | Prior knowledge |
|  | Infinitesimal | - | 0.56 | 0.62 |
|  | Fully connected | 0.15 | - | 0.76 |
|  | Prior knowledge | 0.17 | 0.42 | - |

**Figure S1:**
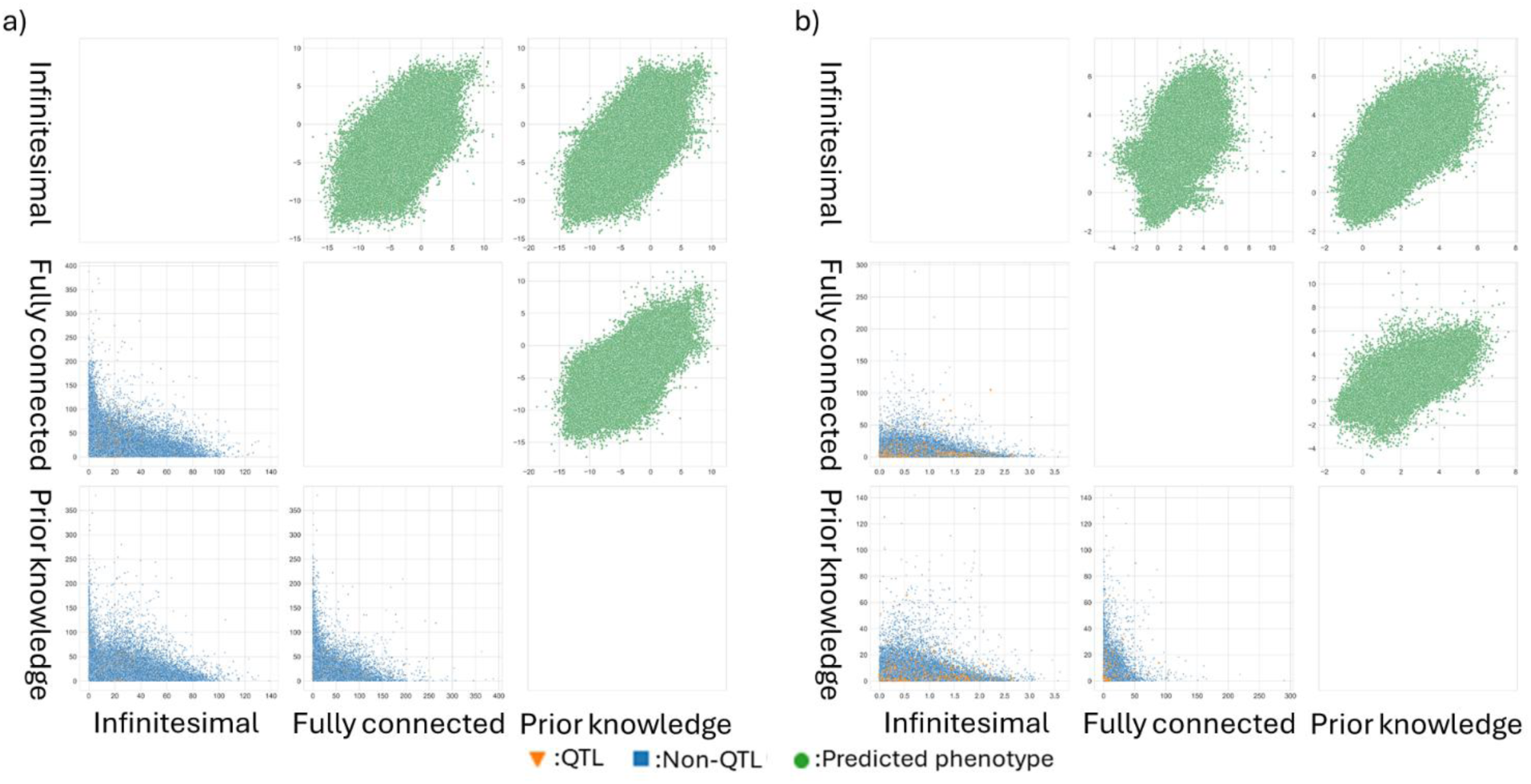
Pairwise comparisons of the GAT models (infinitesimal, fully connected and data-driven prior knowledge) for the anthesis and silking interval (ASI) trait across all the prediction scenarios for a) the TeoNAM (12,500 prediction scenarios) and b) MaizeNAM (6,250 prediction scenarios) datasets at the phenotype and genome levels. Within each subplot in the scatter plot matrix, the GAT models were compared at the predicted phenotypes (top right triangle) and genomic marker effects (the bottom left triangle) levels. The green dots represent a pair of predicted phenotypes for RIL samples in the test sets for each prediction scenario. The blue squares and orange triangles represent pairs of inferred effects of genomic markers in each sample scenario for non-QTL and QTL, respectively. Non-QTL and QTL markers were identified by Chen et al. (2019) for the TeoNAM dataset and Buckler et al. (2009) for the MaizeNAM dataset.

**Figure S2:**
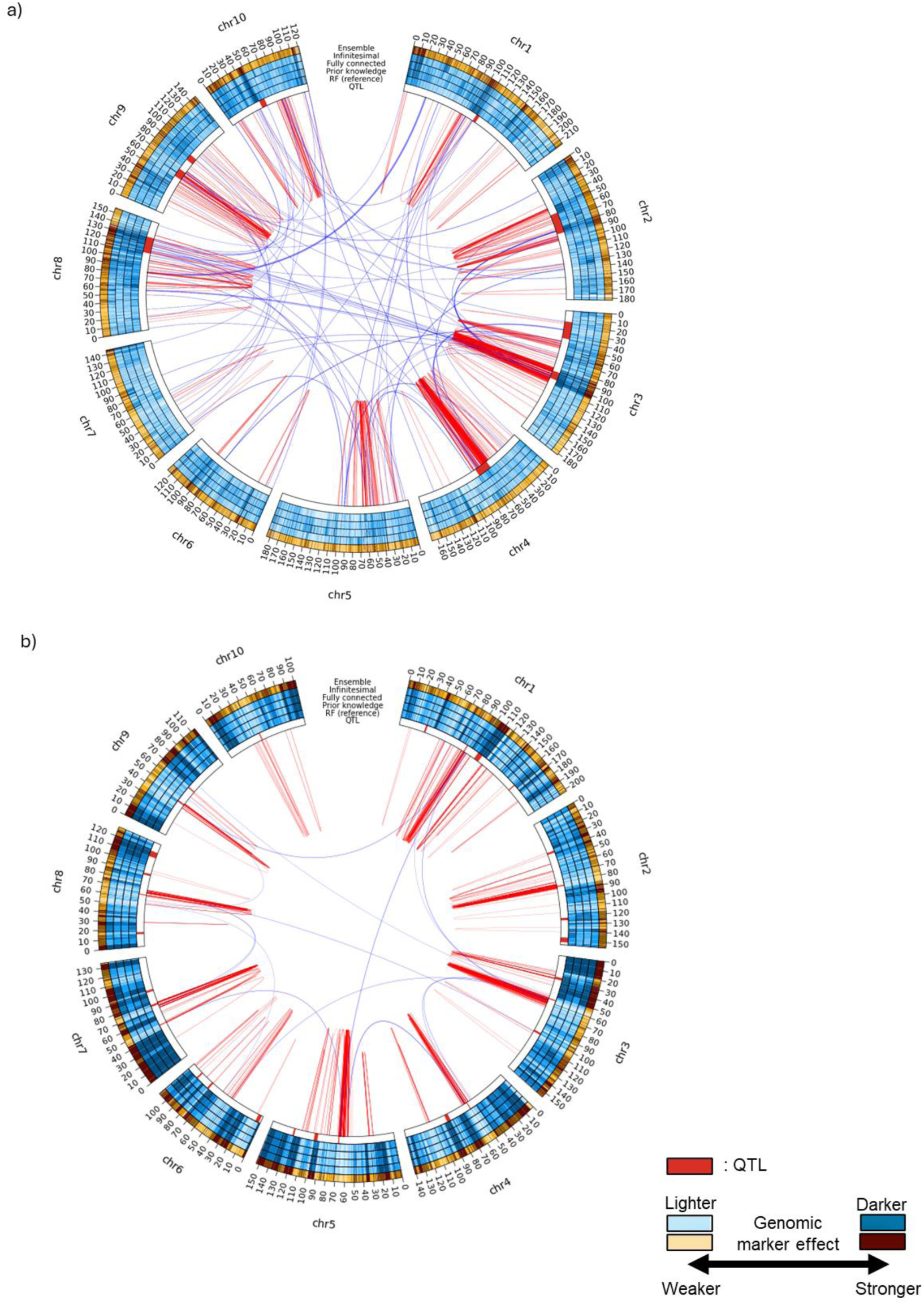
Circos plots for the anthesis to silking interval (ASI) trait for the a) TeoNAM and b) MaizeNAM datasets. The innermost ring (QTL) represents QTL regions in red identified by Chen et al. (2019) for the TeoNAM dataset and Buckler et al. (2009) for the MaizeNAM dataset, respectively. The subsequent four blue rings represent the genomic marker effects from random forest as reference (RF (reference)) used to extract data-driven prior knowledge, the data-driven prior knowledge model, the fully connected model and the infinitesimal model. The outermost ring in orange shows the estimated genomic marker effects of the naïve ensemble-average model. The intensity of the colours shows the strength of the genomic marker effects represented in ten levels based on the quantiles. The colour becomes darker as the genomic marker effects become higher. Links between two genomic marker regions represent genomic marker-by-marker interactions detected by RF using the Shapley scores. The genomic marker pairs with the Shapley scores included in the top 0.001% were selected for the TeoNAM dataset and 0.05% for the MaizeNAM dataset, considering the difference in the total number of identified marker combinations. Thicker lines show stronger marker-by-marker interactions. The red and blue lines represent genomic marker-by-marker interactions within and between chromosomes, respectively. The genetic distance is represented in centimorgans (cM).

**Figure S3:**
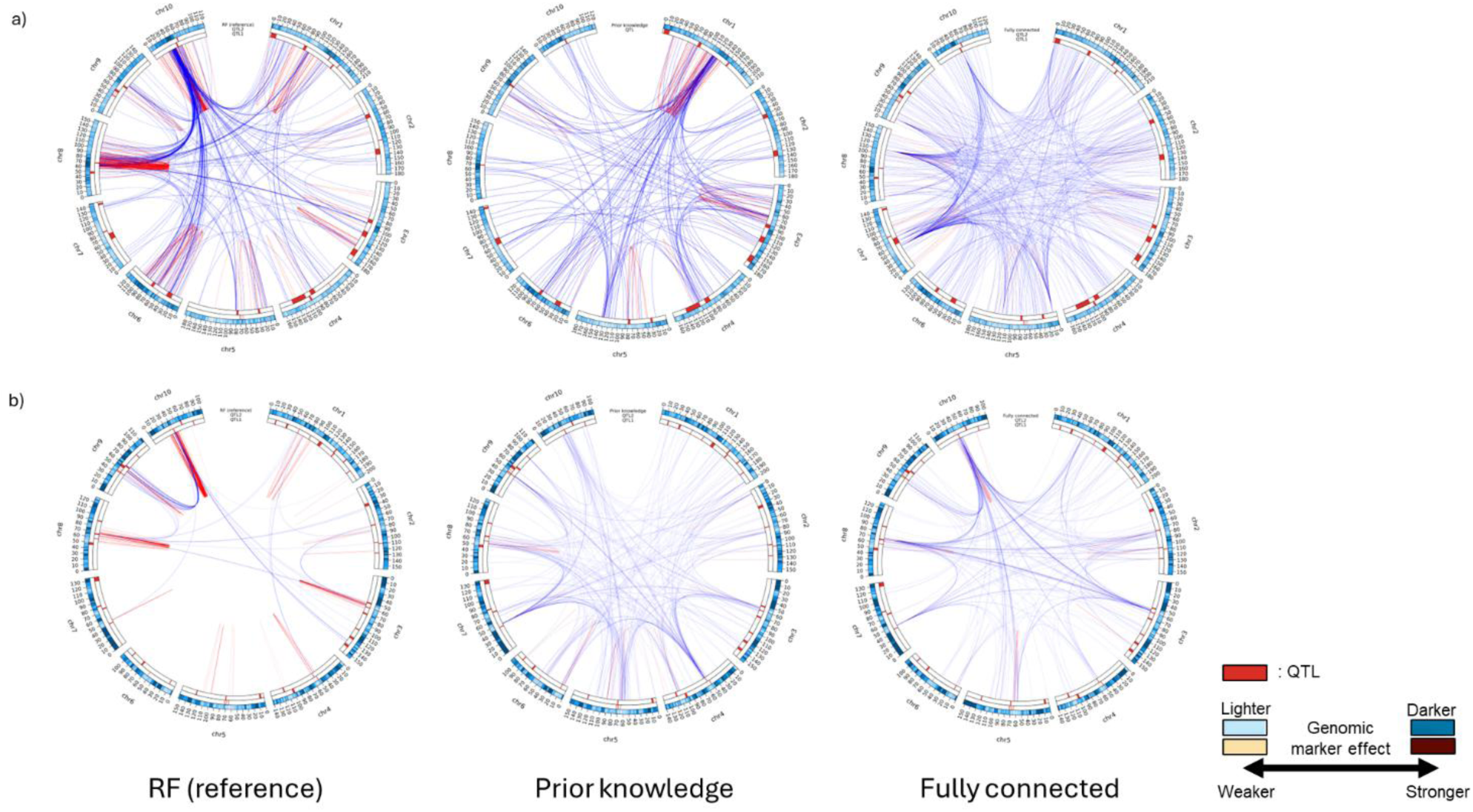
Comparison of inferred gene networks between RF, GAT fully connected and GAT data-driven prior knowledge model for the days to anthesis (DTA) in a) the TeoNAM and b) MaizeNAM datasets. Links between two genomic marker regions represent genomic marker-by-marker interaction effects. For the RF, pairwise Shapley scores were extracted as marker-by-marker interaction effects, while attention values assigned to edges were extracted from the GAT models as marker-by-marker interaction effects. The genomic marker pairs with interaction effects included in the top 0.001% were selected for the TeoNAM dataset and 0.05% for the MaizeNAM dataset, considering the difference in the total number of identified marker combinations. Thicker lines show stronger marker-by-marker interactions. The red and blue lines represent genomic marker-by-marker interactions within and between chromosomes, respectively. The innermost ring (QTL1) represents QTL regions in red identified by Chen et al. (2019) for the TeoNAM dataset and Buckler et al. (2009) for the MaizeNAM dataset, respectively. The second innermost ring (QTL2) represents the QTL regions identified by Wisser et al. (2019). The subsequent blue ring represents the genomic marker effects from each prediction model. The intensity of the colours shows the strength of the genomic marker effects represented in ten levels based on the quantiles. The colour becomes darker as the genomic marker effects become higher. The genetic distance is represented in centimorgans (cM).

**Figure S4:**
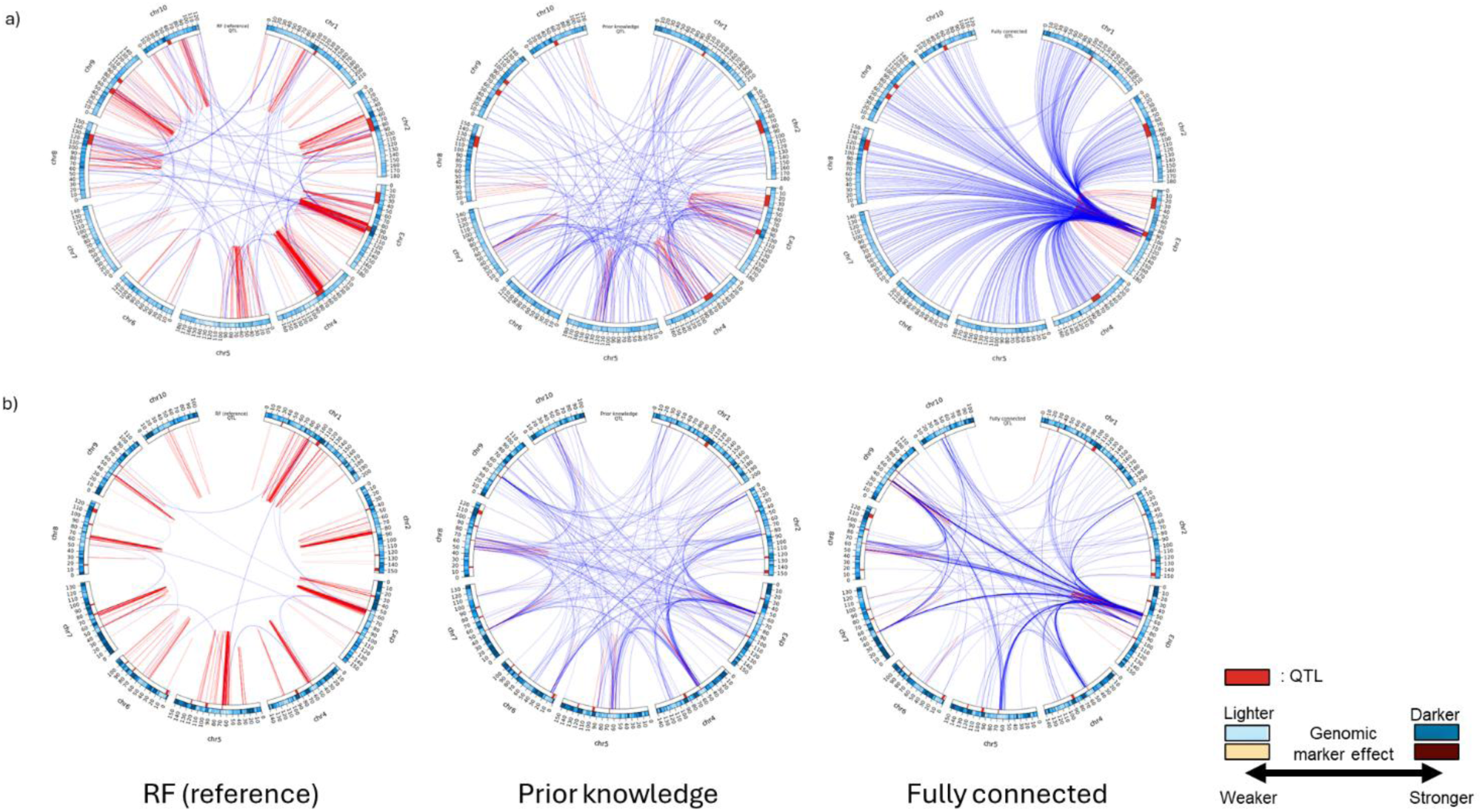
Comparison of inferred gene networks between RF, GAT fully connected and GAT data-driven prior knowledge model for the anthesis to silking interval (ASI) in a) the TeoNAM and b) MaizeNAM datasets. Links between two genomic marker regions represent genomic marker-by-marker interaction effects. For the RF, pairwise Shapley scores were extracted as marker-by-marker interaction effects, while attention values assigned to edges were extracted from the GAT models as marker-by-marker interaction effects. The genomic marker pairs with interaction effects included in the top 0.001% were selected for the TeoNAM dataset and 0.05% for the MaizeNAM dataset, considering the difference in the total number of identified marker combinations. Thicker lines show stronger marker-by-marker interactions. The red and blue lines represent genomic marker-by-marker interactions within and between chromosomes, respectively. The innermost ring (QTL) represents QTL regions in red identified by Chen et al. (2019) for the TeoNAM dataset and Buckler et al. (2009) for the MaizeNAM dataset, respectively. The subsequent blue ring represents the genomic marker effects from each prediction model. The intensity of the colours shows the strength of the genomic marker effects represented in ten levels based on the quantiles. The colour becomes darker as the genomic marker effects become higher. The genetic distance is represented in centimorgans (cM).

